# Chemogenetic stimulation of phrenic motor output and diaphragm activity

**DOI:** 10.1101/2024.04.12.589188

**Authors:** Ethan S. Benevides, Prajwal P. Thakre, Sabhya Rana, Michael D. Sunshine, Victoria N. Jensen, Karim Oweiss, David D. Fuller

## Abstract

Impaired respiratory motor output contributes to morbidity and mortality in many neurodegenerative diseases and neurologic injuries. We investigated if expressing designer receptors exclusively activated by designer drugs (DREADDs) in the mid-cervical spinal cord could effectively stimulate phrenic motor output to increase diaphragm activation. Two primary questions were addressed: 1) does effective DREADD-mediated diaphragm activation require focal expression in phrenic motoneurons (vs. non-specific mid-cervical expression), and 2) can this method produce a sustained increase in inspiratory tidal volume? Wild type (C57/bl6) and ChAT-Cre mice received bilateral intraspinal (C4) injections of an adeno-associated virus (AAV) encoding the hM3D(Gq) excitatory DREADD. In wild-type mice, this produced non-specific DREADD expression throughout the mid-cervical ventral horn. In ChAT-Cre mice, a Cre-dependent viral construct was used to drive neuronal DREADD expression in the C4 ventral horn, targeting phrenic motoneurons. Diaphragm EMG was recorded in isoflurane-anesthetized spontaneously breathing mice at 4-9 weeks post-AAV delivery. The DREADD ligand JHU37160 (J60) caused a bilateral, sustained (>1 hour) increase in inspiratory EMG bursting in both groups; the relative increase was greater in ChAT-Cre mice. Additional experiments in ChAT-Cre rats were conducted to determine if spinal DREADD activation could increase inspiratory tidal volume (VT) during spontaneous breathing, assessed using whole-body plethysmography without anesthesia. Three-to-four months after intraspinal (C4) injection of AAV driving Cre-dependent hM3D(Gq) expression, intravenous J60 resulted in a sustained (>30 min) increase in VT. Subsequently, phrenic nerve recordings performed under urethane anesthesia confirmed that J60 evoked a > 200% increase in inspiratory output. We conclude that targeting mid-cervical spinal DREADD expression to the phrenic motoneuron pool enables ligand-induced, sustained increases in phrenic motor output and VT. Further development of this technology may enable application to clinical conditions associated with impaired diaphragm activation and hypoventilation.

## INTRODUCTION

Many respiratory disorders are associated with reduced or impaired activation of respiratory motoneurons. Neurologic injuries (e.g., traumatic spinal cord injury, stroke) and neurodegenerative conditions (e.g., ALS, Pompe disease) will often result in decreased respiratory motor output, including impaired activation of the phrenic motoneurons which innervate the diaphragm^1–6^. Another prominent example is obstructive sleep apnea, in which pharyngeal motoneurons have reduced output during sleep^7^. Treatment options that increase respiratory motoneuron activation to improve breathing are limited. However, designer receptors exclusively activated by designer drugs (DREADDs) may have use in this regard^8^. Structurally derived from naturally occurring G-protein coupled receptors, DREADDs have been engineered to respond exclusively to exogenous ligands that are otherwise biologically inert^9–11^. This provides a means to selectively stimulate cells expressing the DREADD. Prior studies have used DREADDs to stimulate upper airway muscle activation during breathing^8^. For example, following expression of DREADDs in murine hypoglossal motoneurons, tongue electromyogram (EMG) activity can be increased using DREADD ligands^12–16^. This response is functionally beneficial as shown by increased patency of the upper airway^12^.

The present study focused on chemogenetic activation of the phrenic neuromuscular system. Phrenic motoneurons provide motor innervation of the diaphragm muscle and are located in the mid-cervical (C3-5) spinal cord^17^. We tested the hypothesis that expressing DREADDs in the mid-cervical spinal cord would enable systemic (intravenous or intraperitoneal) delivery of a selective DREADD ligand to produce sustained increases in the respiratory-related activation of the diaphragm. In doing so, we addressed two important questions. First, we determined if effective diaphragm activation requires focal DREADD expression targeting phrenic motoneurons, or if non-specific expression in mid-cervical interneurons and phrenic motoneurons would be sufficient. This question derives from prior studies of cervical spinal cord stimulation. A compelling body of work, with studies in multiple species, demonstrates that non-specific activation of cervical spinal networks can be highly effective at increasing diaphragm activation^18–20^. One theory to explain this result is that a general increase in the excitability of cervical propriospinal networks leads to increased phrenic motoneuron activation^21–23^. There is also evidence that phrenic motoneurons can integrate multiple synaptic inputs in a manner that produces orderly recruitment^18^. Accordingly, DREADD-induced activation of mid-cervical neurons or networks may be sufficient for ligand-induced diaphragm activation. On the other hand, DREADD expression may need to be restricted to phrenic motoneurons if the goal is to produce inspiratory-related diaphragm activation. To address this question, we studied diaphragm responses in two mouse models: 1) a wild-type model in which DREADDs were non-specifically expressed in the C3-5 spinal cord, encompassing interneurons populations and motoneurons, and 2) a choline acetyltransferase (ChAT)-Cre transgenic model in which DREADD expression was restricted to ChAT-positive neurons in the ventral C3-5 spinal cord, targeting the phrenic motoneuron pool.

The second question we addressed was if phrenic motoneuron activation via cervical spinal cord DREADDs could produce a sustained increase in inspiratory tidal volume in unanesthetized, spontaneously breathing animals. While the previous results from the hypoglossal motor system^12,14–16^ provide a proof-of-concept that DREADDs can stimulate respiratory motoneuron activity, whether a sustained increase in tidal volume could be evoked by expressing DREADDs in phrenic motoneuron was uncertain. For example, during spontaneous breathing, a DREADD-induced increase in phrenic motoneuron excitability, and thus diaphragm activation, could be rapidly offset by decreases in bulbospinal neural drive to the phrenic motor pool, secondary to reduced arterial CO_2_ or increased vagal-mediated inhibition. An increase in diaphragm activation could also trigger a decrease in accessory respiratory muscle activation, thereby attenuating or preventing increases in tidal volume. Lastly, data from the hypoglossal motor system^12,14–16^, as well as our initial results in the anesthetized mouse indicated that both phasic (i.e., during the inspiratory period) and tonic (i.e., occurring across the respiratory cycle) activation of the diaphragm would increase after DREADD activation, and how this would impact tidal volume was not clear. Accordingly, we studied ChAT-Cre rats using whole-body plethysmography and a direct measure of phrenic motor output via nerve recordings. The plethysmography studies allowed us to determine if DREADD activation of phrenic motoneurons causes a sustained increase in tidal volume and ventilation during spontaneous breathing in the unanesthetized rat. The nerve recordings were done under urethane anesthesia and enabled direct quantification of DREADD activation on the neural drive of the diaphragm while controlling variables including arterial CO_2_ and lung volume.

Collectively the results of this work demonstrate that mid-cervical spinal DREADD expression enables the J60 ligand to produce a sustained increase in the neural drive to the diaphragm, producing an increase in tidal volume during spontaneous breathing. Further development of this technology may enable application to clinical conditions associated with impaired diaphragm activation and hypoventilation.

## RESULTS

### Diaphragm EMG responses in wild-type mice

Wild-type mice underwent bilateral injections of AAV9-hSyn-HA-hM3D(Gq)-mCherry into the ventral horns at spinal segment C4. Following a four-to-five-weeks incubation period, mice underwent terminal diaphragm EMG recordings before and after application of the selective DREADD ligand, J60. On average, wild-type mice showed increases in diaphragm EMG output in at least one hemidiaphragm after J60 administration (**Figure 1**). The area under the curve (AUC) of the rectified and integrated diaphragm EMG was significantly increased after DREADD activation (**Figure 1d**) in both the left (**p = 0.002**; **Table S1**) and right (**p = 0.002**; **Table S1**) hemidiaphragm. Additionally, the peak-to-peak amplitude of the rectified and integrated diaphragm EMG burst activity increased bilaterally following J60 administration (<u>Left hemidiaphragm</u>: p = 0.056; <u>Right</u> <u>hemidiaphragm</u>: **p = 0.01**; **Figure 1e**; **Table S1**). Lastly, the tonic activity of the diaphragm was assessed (**Figure 1f**). Similar to the previous measures of EMG output both the left (**p = 0.052**; **Table S1**) and right (**p < 0.001**; **Table S1**) hemidiaphragm exhibited a significant increase in tonic activity following J60 administration. Respiratory rate was consistent for the duration of the experiment. Notably, there was no substantial change in the respiratory rate of these spontaneous breathing mice after J60 administration (p = 0.863; **Figure 1g**; **Table S1**).

**Figure 1.**
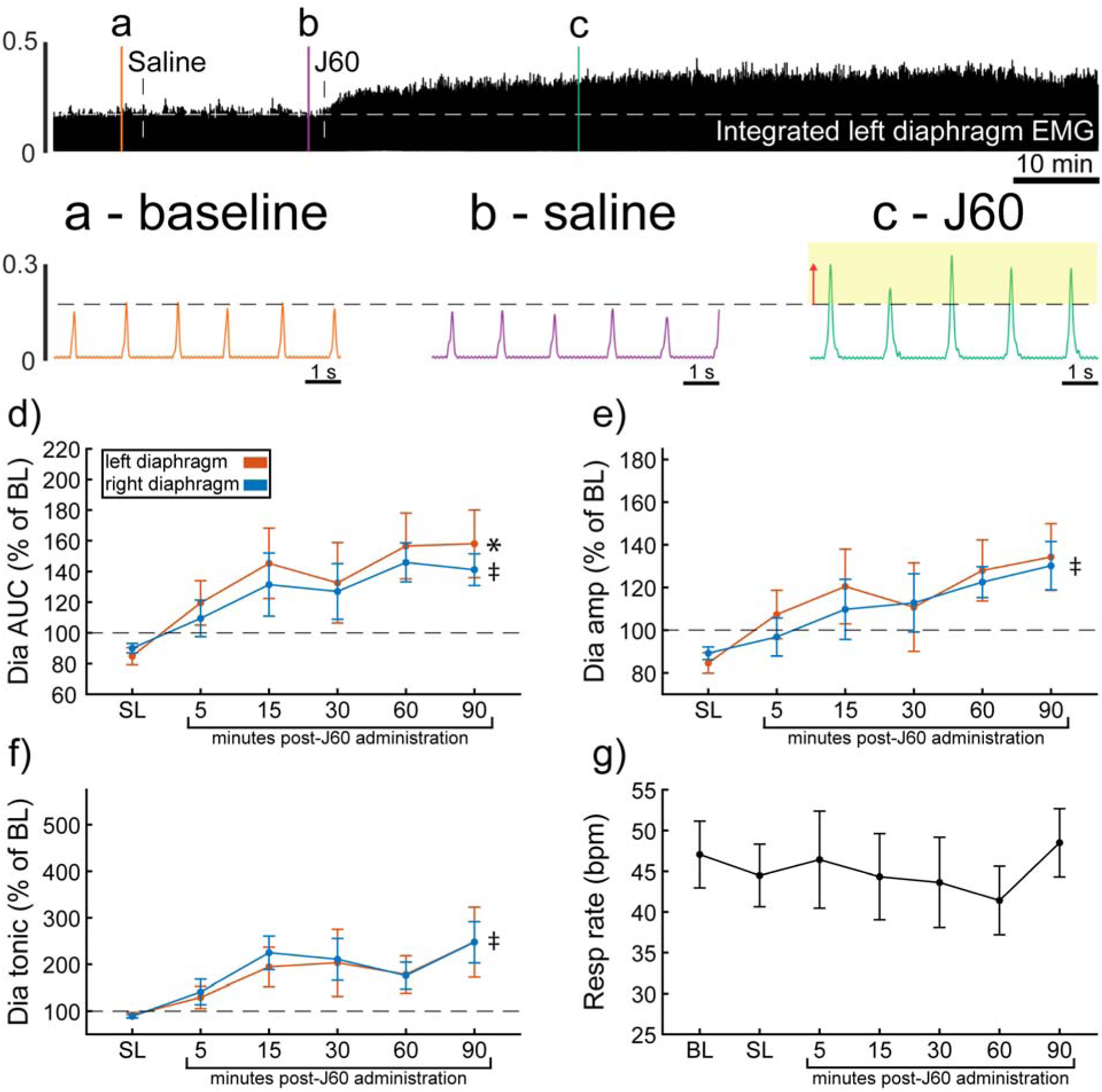
DREADD activation increases diaphragm EMG output in wild-type mice. A representative example of diaphragm EMG activity before and after application of the J60 DREADD ligand is shown in the top panel. Examples of the individual inspiratory EMG bursts at baseline (a), after vehicle (b), and after J60 (c) are shown. The J60 ligand increased diaphragm output but did not impact respiratory rate. The mean responses (n = 11; n = 7 females) for EMG AUC, peak-to-peak amplitude, tonic activity, and respiratory rate are shown in panels d-g. For diaphragm EMG data (panels d-f) left hemidiaphragm EMG is represented in orange, while right hemidiaphragm EMG is blue. Error bars depict ±D1 SEM. Statistical reports for all panels are provided in Supplemental Table 1. * and ‡ symbols indicate significant main effects (p < 0.05) on One-Way RM ANOVA for the left and right hemidiaphragm, respectively. Dia = diaphragm, AUC = area under the curve, amp = peak amplitude, BL = baseline, SL = saline (sham injection).

In all experiments, the selective DREADD ligand, J60, produced an increase in diaphragm EMG burst amplitude during inspiration. However, this increase was not always detected in both the left and right hemidiaphragm EMG recordings. Five mice showed a bilateral increase in diaphragm output after J60 administration (**Figure 1**), four mice showed a response that was limited to the right hemidiaphragm, and two showed a response that was limited to the left hemidiaphragm (**Figure 1**).

### Diaphragm EMG responses in ChAT-Cre mice

ChAT-Cre mice received bilateral intraspinal injections of AAV9-hSyn-DIO-hM3D(Gq)-mCherry into the ventral horns at C4. ChAT-Cre mice underwent terminal diaphragm EMG recording using the same protocol as wild-type mice, following a four-to-nine-week incubation. All mice (n = 9/9) showed an increase in diaphragm EMG output in response to the J60 DREADD ligand in at least one hemidiaphragm (**Figure 2**). On average, both left (**p = 0.011**; **Table S2**) and right (**p < 0.001**; **Table S2**) diaphragm EMG AUC increased over time after J60 administration (**Figure 2d**). Diaphragm EMG peak-to-peak amplitude had a similar, bilateral increase following J60 delivery (<u>Left hemidiaphragm</u>: **p = 0.013**; <u>Right hemidiaphragm</u>: **p < 0.001**; **Table S2**; **Figure 2e**). Lastly, tonic activity also showed an increase over time after J60 administration (**Figure 2f**) for both the left (**p = 0.002**; **Table S2**) and right (**p < 0.001**; **Table S2**) hemidiaphragm. Respiratory rate decreased significantly over time after J60 administration (**p < 0.001**; **Figure 2g**; **Table S2**). Apart from two mice that showed a unilateral EMG response that was limited to the right hemidiaphragm the remaining ChAT-Cre mice (n = 7/9) had bilateral increases in diaphragm EMG output following J60 administration (**Figure 2**).

**Figure 2.**
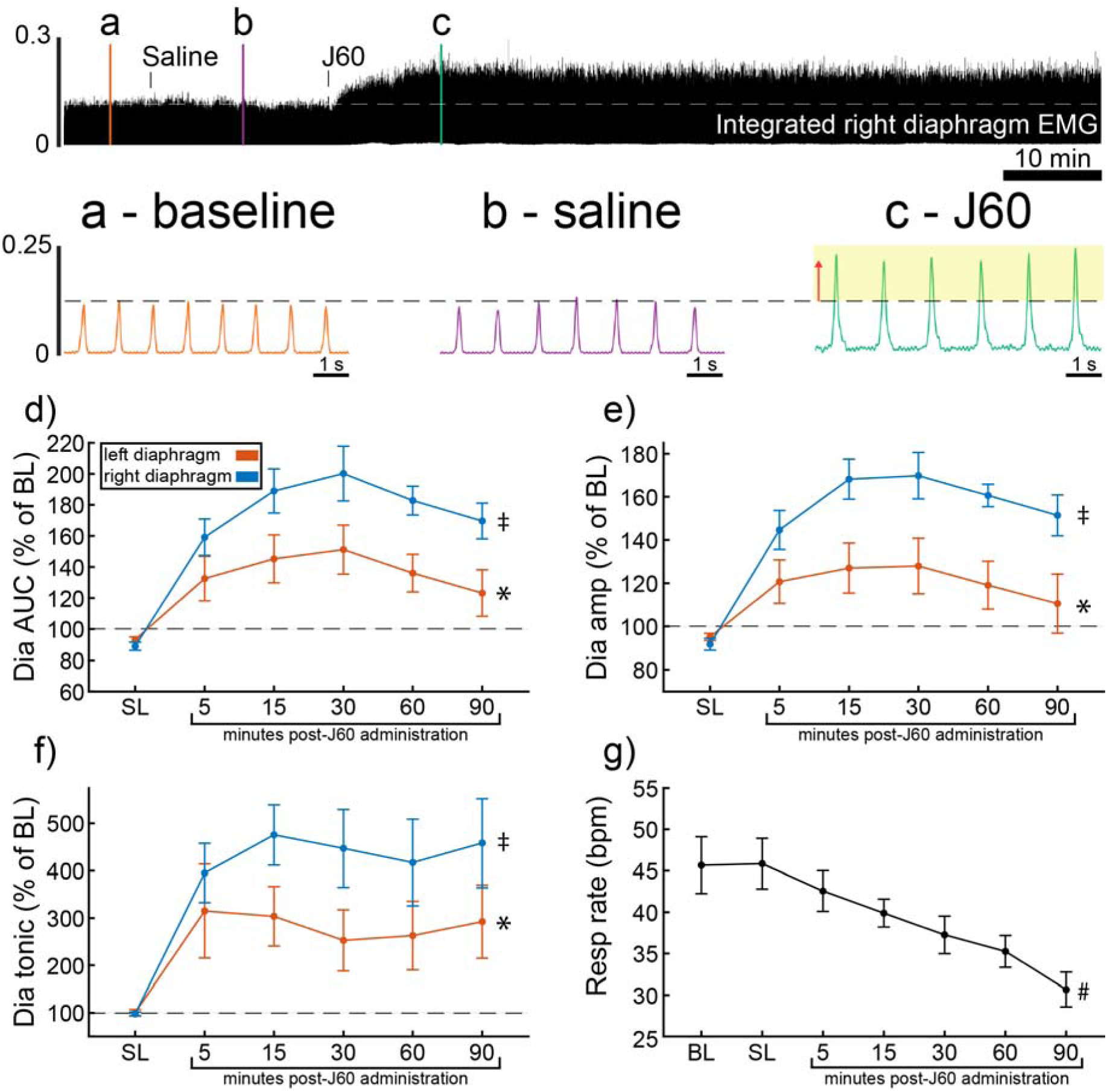
DREADD activation increases diaphragm EMG output in ChAT-Cre mice. A representative example of diaphragm EMG activity before and after application of the J60 DREADD ligand is shown in the top panel. Examples of the individual inspiratory EMG bursts at baseline (a), after vehicle (b), and after J60 (c) are shown. Mean responses (n = 9; n = 6 females) for EMG AUC, peak-to-peak amplitude, tonic activity, and respiratory rate are shown in panels d-g. The DREADD ligand caused a bilateral increase in diaphragm EMG AUC, peak-to-peak amplitude, and tonic activity. For all EMG parameters, the responses were greater on the right vs. left hemidiaphragm. Respiratory rate decreased over time. For panels d-f, the left hemidiaphragm EMG is represented in orange, while right hemidiaphragm EMG is blue. Error bars depict ±D1 SEM. Statistical reports for all panels are provided in Supplemental Table 2. * and ‡ symbols indicate significant main effects (p < 0.05) on One-Way RM ANOVA for the left and right hemidiaphragm, respectively. # indicates a significant main effect (p < 0.05) on One-Way RM ANOVA for respiratory rate data. Dia = diaphragm, AUC = area under the curve, amp = peak amplitude, BL = baseline, SL = saline (sham injection).

### Wild type vs. ChAT-Cre comparison

Diaphragm EMG responses of wild-type and ChAT-Cre mice were compared at the 30-minute post-J60 time point (**Figure 3**). Left hemidiaphragm responses to J60 were similar between the two groups across all outcome measures (<u>AUC</u>: p = 0.998; <u>Peak-to-peak amplitude</u>: p = 0.771; <u>Tonic activity</u>: p = 0.160; **Table S3**; **Figure 3a-c**). However, right hemidiaphragm responses to J60 differed across AUC (**Figure 3d**), peak-to-peak amplitude (**Figure 3e**), and tonic activity (**Figure 3f**) with ChAT-Cre mice on average having larger magnitude responses compared to wild-type mice (<u>AUC</u>: **p = 0.0417**; <u>Peak-to-peak amplitude</u>: **p = 0.00403**; <u>Tonic activity</u>: **p = 0.00207**; **Table S3**). Respiratory rate was not different between the two groups (p = 0.382; **Table S3**; **Figure 3g**).

**Figure 3.**
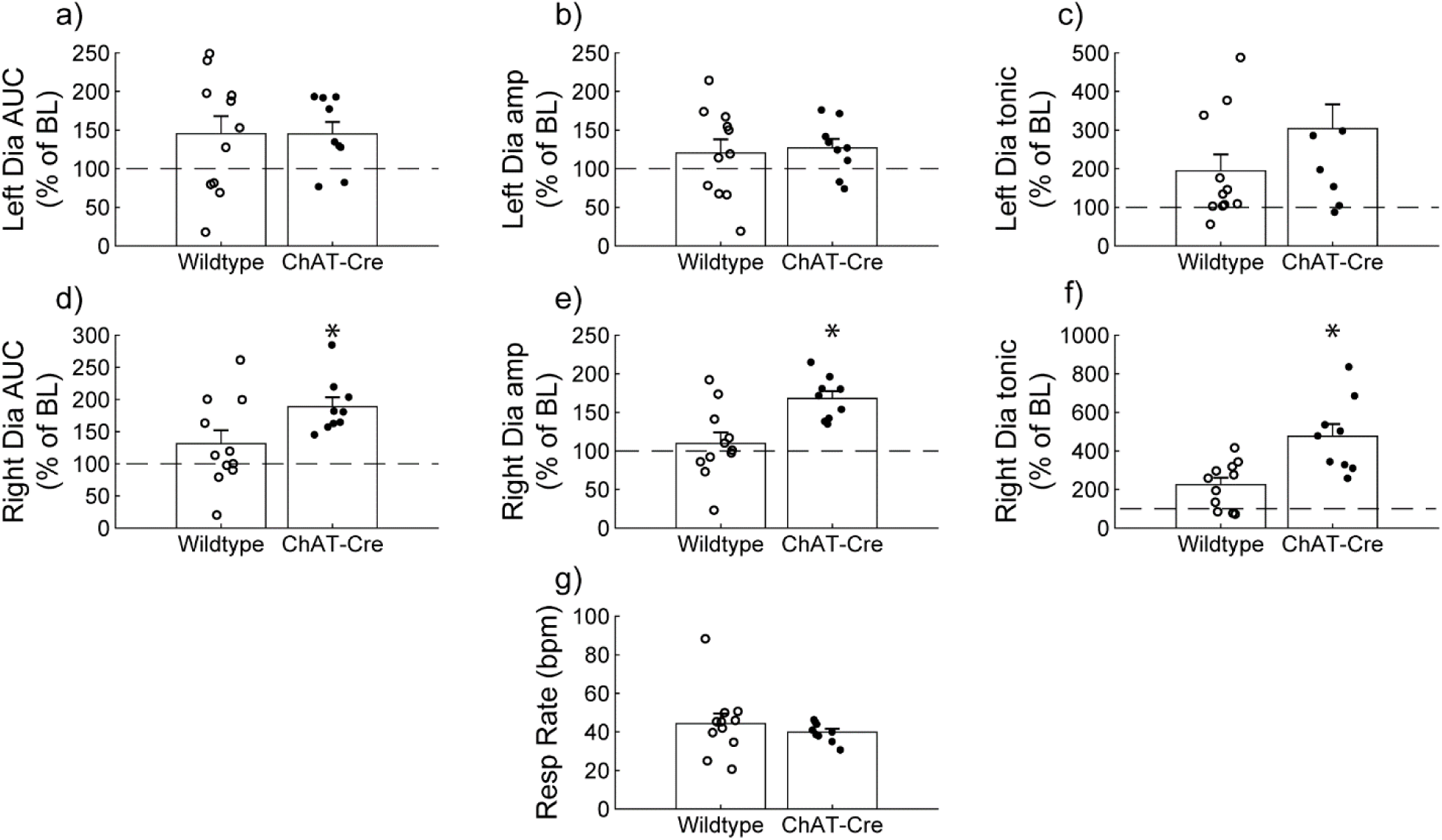
Wild type vs. ChAT-Cre mouse responses to DREADD activation. Direct comparisons of diaphragm EMG response parameters (a-f) and respiratory rate (g) at 30-minute post-J60 application (Wild type, n = 11; n = 7 females; ChAT-Cre, n = 9; n = 6 females). Left hemidiaphragm EMG AUC (a), peak-to-peak amplitude (b), and tonic activity (c) were similar between groups. However, the same parameters on the right hemidiaphragm (d-f) were greater in ChAT-Cre mice. Respiratory rate was similar between groups. Error bars depict ±ℒ1 SEM. Statistical reports for all panels are provided in Supplemental Table 3. * p < 0.05. AUC = area under the curve, amp = peak EMG amplitude, Dia = diaphragm, BL = baseline, resp rate = respiratory rate.

The *a priori* expected recording duration for these experiments in anesthetized and spontaneously breathing mice was 90-minute. However, five of the eleven total wild-type mice in this experiment did not survive for this duration. It is unclear if this was a non-specific result associated with prolonged anesthesia, or if this was physiologically related to DREADD activation. No mice had evidence of adverse reaction in the initial 30-minutes following delivery of J60. Of the five mice which did not survive, three mice died between the 30- and 60-minute time points after J60, and two mice died just prior to the 90-minute time point. In contrast, all mice in the ChAT-Cre cohort (9 of 9) survived the total duration of the experimental protocol. A chi-square evaluation of the survival proportions was not statistically significant (Chi-squared = 3.2997, df = 1, p = 0.06929). However, considering the sample size, the results suggest some association between mouse strain (i.e., wild type, ChAT-Cre) and death, suggesting that non-specific DREADD activation may be contraindicated.

### J60 control experiments

The DREADD ligand was administered to wild-type animals with no hM3D(Gq) expression in the mid-cervical spinal cord (**Figure S1**). This was done to determine the impact of J60 administration on diaphragm EMG in the absence of DREADD expression (n= 2 C57/bl mice; n= 3 Sprague Dawley rats). There was no discernable impact of J60 on the diaphragm EMG burst amplitude (mV) (**Figure S1a-b**). Responses were also not different between sham (saline) and J60 when normalized to baseline activity (**Figure S1c-d**).

### ChAT-Cre rats – Plethysmography and Phrenic Nerve Recordings

A small cohort of ChAT-Cre rats underwent anesthetized diaphragm EMG recordings to ensure DREADD responses similar to the mouse cohorts could be obtained in rats. ChAT-Cre rats (n = 4) underwent bilateral, intraspinal injections of AAV9-hSyn-DIO-hM3D(Gq)-mCherry into the ventral horns at C4 to introduce the hM3D(Gq) DREADD transgene into phrenic motoneurons. Four of four rats showed increased diaphragm EMG output after DREADD activation (**Figure S2**). With that knowledge, we used a separate group of ChAT-Cre rats (n = 9; n = 3 females) to assess the effects of DREADD activation on ventilation. Whole-body plethysmography was used to measure breathing frequency, tidal volume, and minute ventilation before and after intravenous delivery of saline (sham) and J60 (**Figure 4**). Delivery of the J60 ligand resulted in an increase in inspiratory tidal volume compared to sham infusion (<u>Normalized to Weight (ml/kg):</u> Main effect of Treatment: **p = 0.037**; **Figure 4a**; <u>Normalized to Baseline</u>: Main effect of Treatment: p = 0.091; **Figure 4d**; **Table S4**). Respiratory rate appeared to be unaffected by DREADD activation and was similar between sham and J60 conditions (<u>Respiratory Rate</u>: Main effect of Treatment: p = 0.582; **Figure 4b**; <u>Respiratory Rate Normalized to Baseline</u>: Main effect of Treatment: p = 0.774; **Figure 4e**; **Table S4**). Minute ventilation was slightly elevated in the J60 condition vs sham; however, this increase did not reach the threshold for statistical significance (<u>Normalized to Body Weight</u>: Main effect of Treatment: p = 0.194; **Figure 4c**; <u>Normalized to Baseline</u>: Main effect of Treatment: p = 0.337; **Figure 4f; Table S4**). Responses to a hypercapnic-hypoxia ventilatory challenge were also assessed (**Figure S3**). Tidal volume (**Figure S3a**; <u>Normalized to Weight (ml/kg)</u>: p = 0.845; **Figure S3d**; <u>Normalized to baseline</u>: p = 0.643), respiratory rate (**Figure S3b**; <u>Respiratory Rate</u>: p = 0.262; **Figure S3e**; <u>Rate Normalized to Baseline</u>: p = 0.734), and minute ventilation (**Figure S3c**; <u>Normalized to Body Weight</u>: p = 0.697; **Figure S3f**; <u>Rate</u> Normalized to Baseline: p = 0.912) did not differ between J60 vs sham condition during hypercapnic-hypoxic ventilatory challenges.

**Figure 4.**
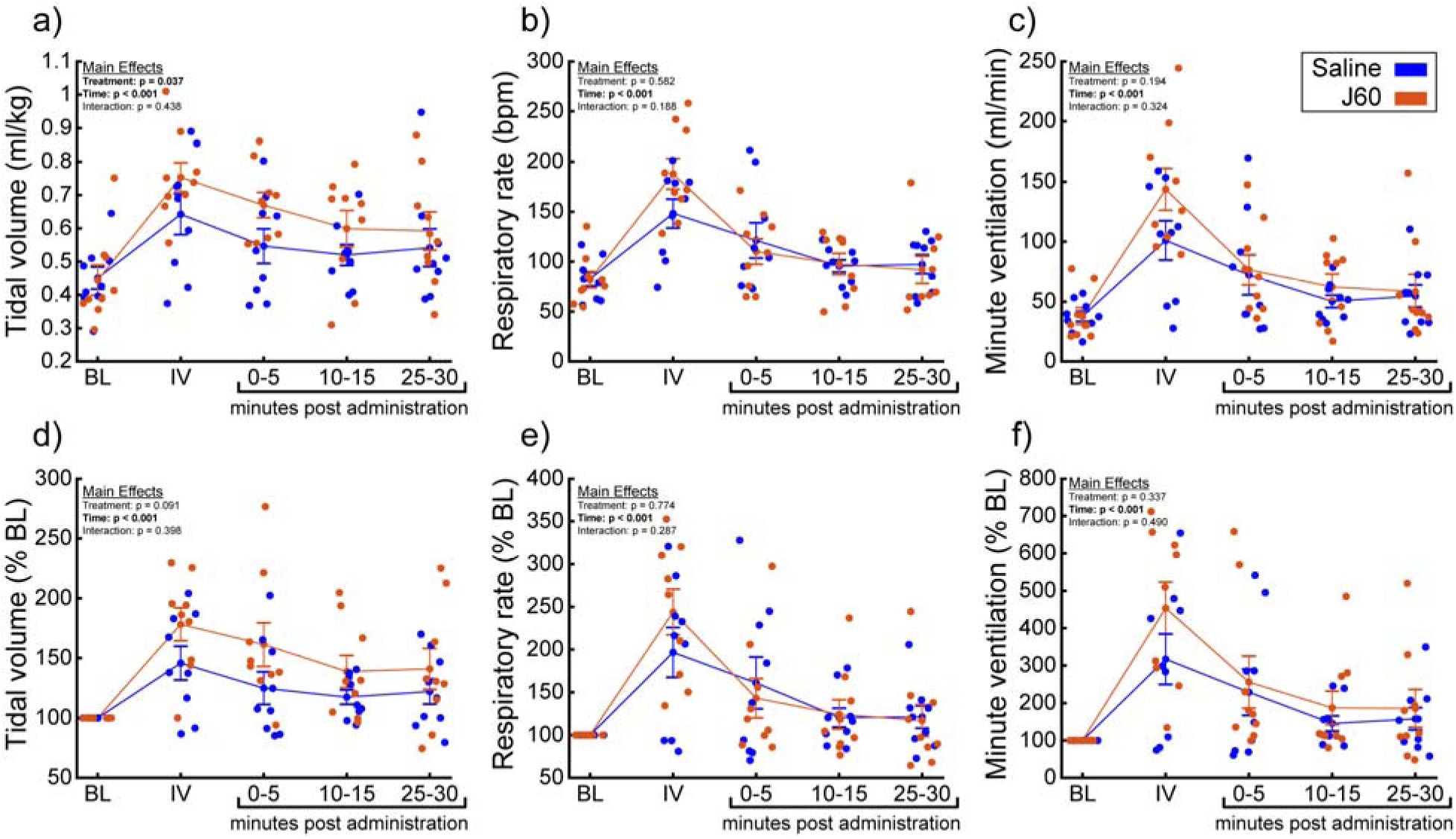
DREADD activation increases ventilation in unanesthetized ChAT-Cre rats. Summary plots (n = 9; n = 3 females) showing the impact of the J60 DREADD ligand on tidal volume, respiratory rate, and minute ventilation are shown in panels a-c. The normalized values (% of baseline) are shown in panels d-f. The DREADD ligand increased tidal volume compared to sham infusion (saline). Error bars depict ±D1 SEM. Statistical reports for all panels are provided in Supplemental Table 4. BL = baseline, IV = intravenous infusion period.

Phrenic nerve recordings were made to directly assess the effects of DREADD activation on phrenic output. There was no detectable relationship between time post-AAV injection and phrenic response to DREADD activation (Pearson correlation; <u>Left peak-to-peak response</u>: p = 0.215; <u>Right peak-</u> <u>to-peak response</u>: p = 0.318).

Application of the J60 ligand caused a rapid, sustained, and bilateral increase in phrenic nerve efferent burst amplitude (<u>Left phrenic peak-to-peak amplitude (normalized to baseline)</u>: **p < 0.001**; <u>Right</u> <u>phrenic peak-to-peak amplitude (normalized to baseline)</u>: **p < 0.001**; **Table S5**; **Figure 5b-c**), whereas saline injection had no impact. The increase in phrenic burst amplitude lasted up to 100 minutes post-J60 administration, at which point the experiment was terminated. Application of the J60 ligand also resulted in an increase in phrenic tonic activity (<u>Left phrenic tonic activity (normalized to baseline)</u>: **p < 0.001**; <u>Right phrenic tonic activity (normalized to baseline)</u>: **p < 0.001**; **Table S5**; **Figure 5d-e**).

**Figure 5.**
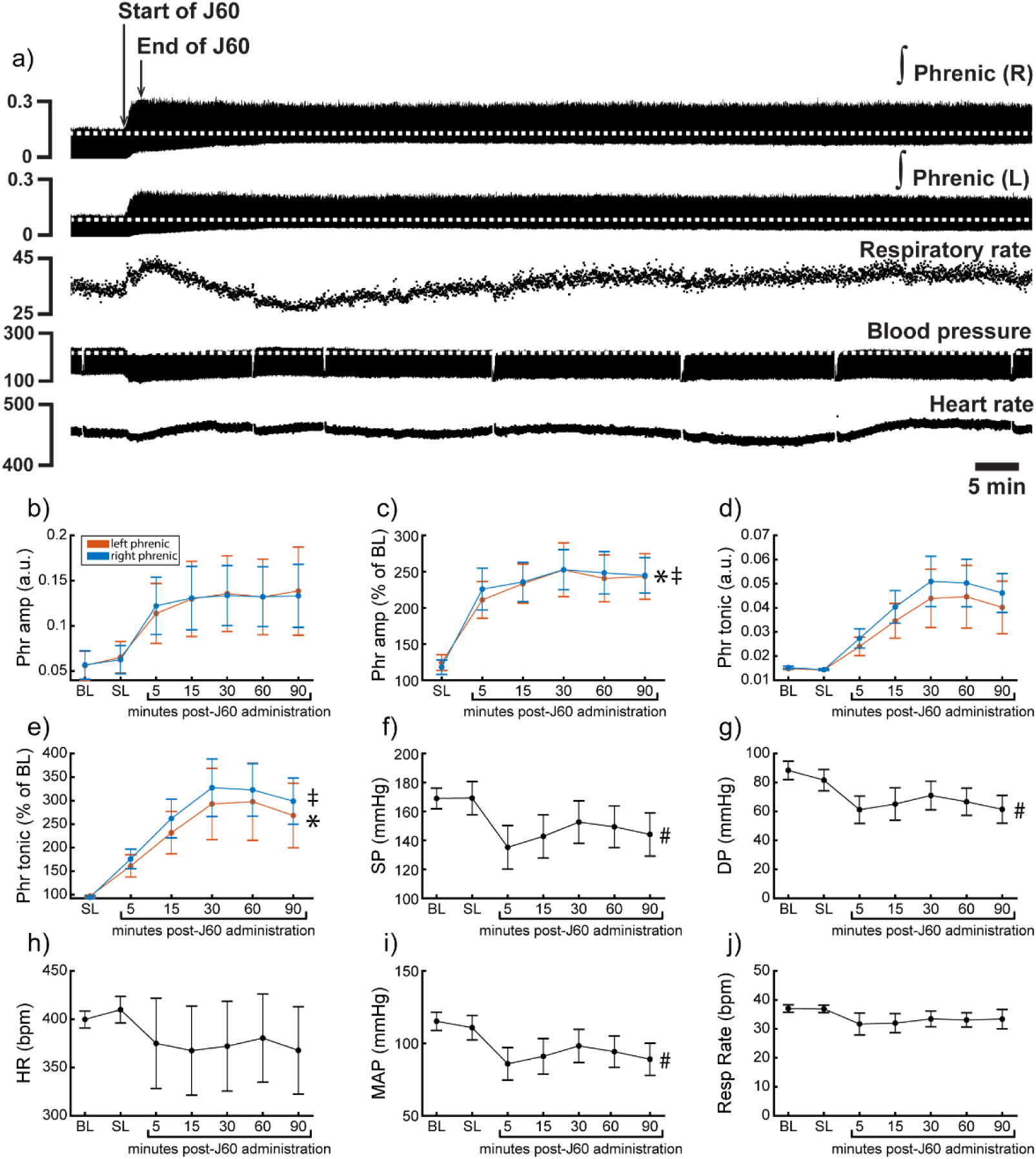
DREADD activation increases phrenic nerve output in ChAT-Cre rats. Representative data showing that the J60 DREADD ligand causes a rapid increase in phrenic nerve output (a). Mean data (n = 9; n = 3 females) showing the impact of J60 application on phrenic nerve raw (b) and normalized (c) peak-to-peak amplitude, raw (d) and normalized (e) tonic activity, systolic blood pressure (f), diastolic blood pressure (g), heart rate (h), mean arterial blood pressure (i), and respiratory rate (j). The J60 ligand caused an increase in phrenic peak-to-peak amplitude and tonic activity. Systolic, diastolic, and mean arterial blood pressure all decreased after J60 application. Heart rate and respiratory rate were not statistically different after J60. In panels b-e, the left phrenic is represented in orange, while right phrenic is blue. Error bars depict ±D1 SEM. Statistical reports for all panels are provided in Supplemental Table 6. * and ‡ symbols indicate significant main effects (p < 0.05) on One-Way RM ANOVA for the left and right hemidiaphragm, respectively. # indicates a significant (p < 0.05) effect on One-Way RM ANOVA for respiratory rate data. Phr = phrenic, amp = amplitude, BL = baseline, SP = systolic pressure, DP = diastolic pressure, HR = heart rate, MAP = mean arterial pressure.

Heart rate, systolic and diastolic blood pressure, mean arterial blood pressure (MAP), as well as respiratory rate, were also assessed (**Figure 5f-j**). Application of J60 did not affect heart rate (p = 0.587; **Table S5**; **Figure 5h**) or respiratory rate (p = 0.282; **Table S5; Figure 5j**) but did result in a decrease in both systolic (**p < 0.001**; **Table S5**; **Figure 5f**) and diastolic blood pressure (**p < 0.001**; **Table S5**; **Figure 5g**) as well as MAP (**p < 0.001**; **Table S5**; **Figure 5i**).

### Histological analysis

We performed a qualitative analysis of the mid-cervical spinal cord from each animal to assess the extent of mCherry fluorophore expression (**Figure S4**). All mice from both cohorts showed evidence of mCherry expression in at least one segment of the mid-cervical spinal cord (**Figure 6**) with the exception n = 1 ChAT-Cre mouse. This mouse was excluded from all analyses based on *a priori* exclusion criteria, which stipulated animals must show evidence of mCherry expression in the grey matter of at least one spinal segment from C3-C6 to be included in the final analysis. A summary of the results is given in **Table 1**.

**Figure 6.**
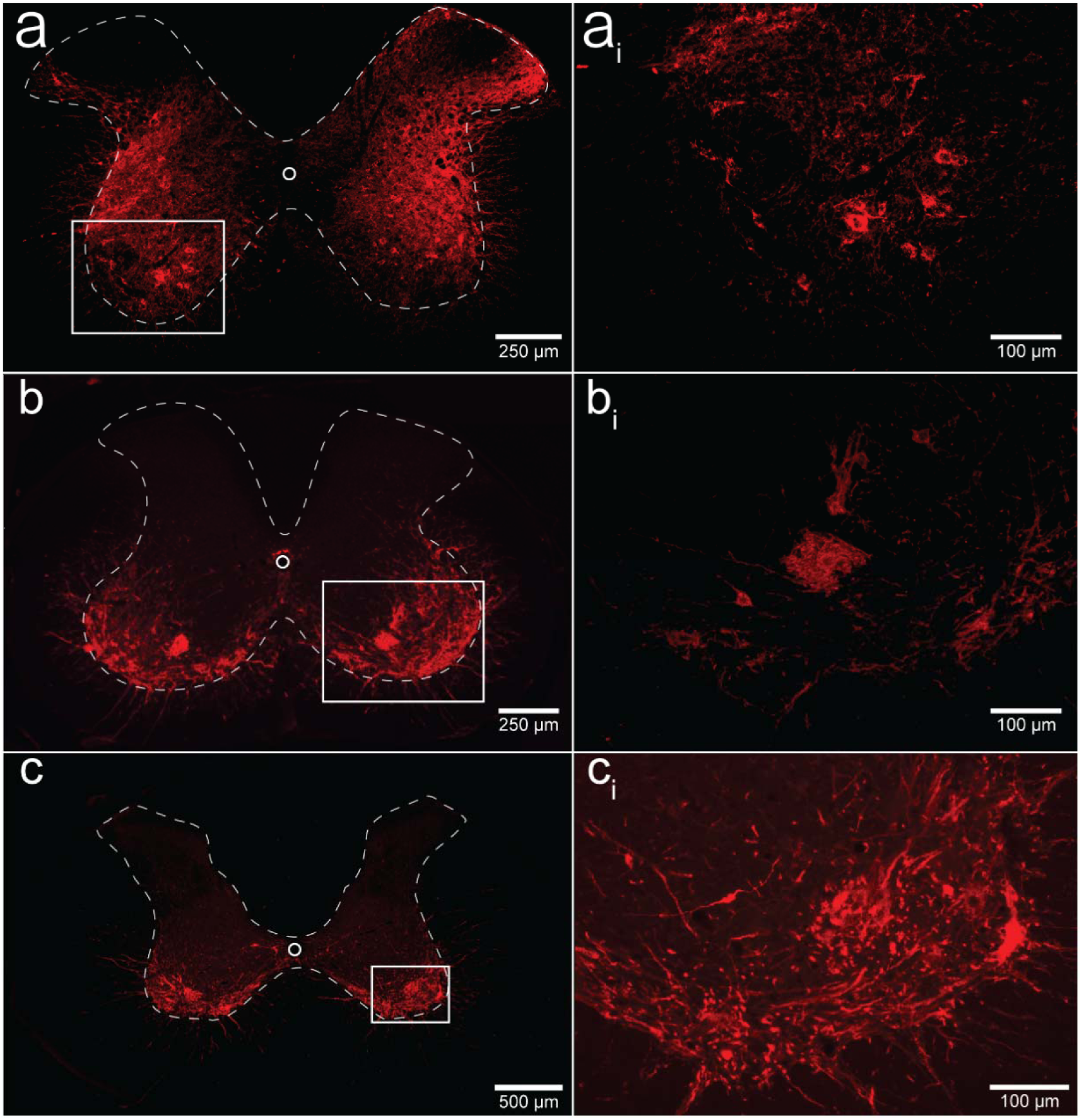
Histological assessment of mCherry expression in the C4/C5 spinal segments. Representative photomicrographs of mid-cervical spinal sections from a wild-type mouse (a-a_i_), a ChAT-Cre mouse (b-b_i_), and a ChAT-Cre rat (c-c_i_). Wild-type mice (a-a_i_) showed a nonspecific pattern of expression throughout the mid-cervical grey matter. ChAT-Cre mice and rats (b-c_i_) showed expression limited to neurons in the ventral horns. Red color indicates positive and mCherry fluorescence. Dashed white line indicates the approximate white-gray matter demarcation.

**Table 1.**
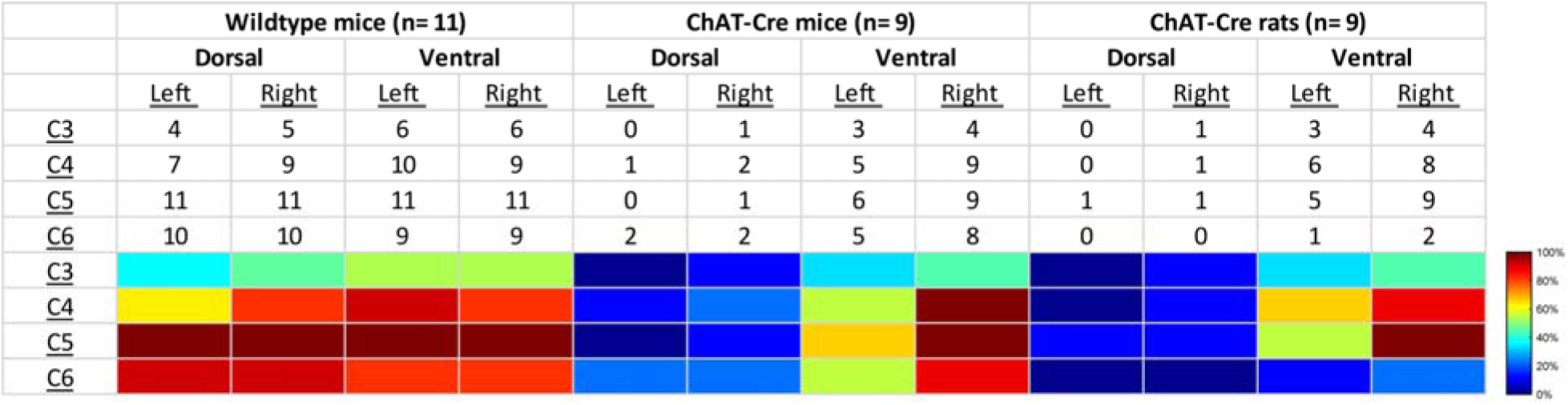
Qualitative assessment of mCherry expression in the mid-cervical spinal cord. Spinal segments C3-C6 were assessed in quadrants broken into dorsal, ventral, left, and right. Spinal segments were counted as “positive” if they showed any evidence of mCherry expression in neuronal soma or fibers. The counts therefore indicate the number of animals of a given cohort that were mCherry positive for a given spinal segment quadrant. All animals showed a slight trend for more mCherry expression moving rostral to caudal and for more expression in the ventral vs the dorsal lamina. This trend was more prominent in the ChAT-Cre animals. At the bottom of the table, a heatmap is provided for easier assessment of the distribution of positive mCherry counts across quadrants and spinal segments.

Patterns of expression were relatively homogenous in wild-type animals. In this cohort, the number of mice with positive mCherry expression in the grey matter increased on the rostral-caudal axis. Positive mCherry counts were comparable on both the dorsal-ventral and left-right axes, with a majority of mice expressing mCherry in all four quadrants. The diaphragm EMG responses to J60, on average, exhibited similarity between the left and right hemidiaphragm in these mice, aligning with the observed pattern of mCherry expression. (**Figure 1d-f**).

In contrast, mCherry expression in the ChAT-Cre mice cohort was more prevalent in the ventral horns and the right side of the cord. Like the wild-type mice, there was a slight trend for increased mCherry expression moving rostral to caudal. Clear mCherry expression was detectable in the spinal cord in of all nine ChAT-Cre mice included in the final data set. One additional ChAT-Cre mouse was excluded from analysis as it showed no evidence of mCherry in the mid-cervical spine. Interestingly, this particular mouse appeared to show a modest increase in diaphragm output in response to the J60 ligand in the left-hemidiaphragm only (∼45% increase compared to baseline activity). While this animal was ultimately excluded from our analysis, it is possible that this mouse did express hM3D(Gq) in the mid-cervical spinal cord but an issue in tissue processing resulted in an inability to visualize the mCherry fluorophore in the spinal tissue. All other ChAT-Cre mice showed robust mCherry expression in the ventral horns of at least one spinal segment from C3-C6. These mice demonstrated a larger average DREADD response in the right hemidiaphragm than the left (**Figure 2d-f**), possibly stemming from the fact that a greater number of mice exhibited mCherry expression on the right side compared to the left (**Figure S5**).

ChAT-Cre rats showed expression predominately in the ventral horns throughout the mid-cervical spinal cords with the highest levels of expression in spinal segments C4 and C5. Expression in this cohort was slightly more prominent on the right side of the cord and in the ventral horn. These histological findings were consistent with the physiological results. Although the magnitude of DREADD response between the left and right phrenic nerves for this cohort was not statistically different, there was a trend of slightly higher right phrenic tonic response compared to the left (**Figure 5b-e**). This trend is mirrored in the pattern of mCherry expression, where expression levels were approximately equal between spinal segments but tended to be slightly higher in the right ventral horns compared to the left.

## DISCUSSION

We describe a novel method to increase diaphragm EMG output by expressing the excitatory DREADD, hM3D(Gq), in the mid-cervical spinal cord, targeting phrenic motoneurons. Following AAV-driven expression of the DREADD in the spinal cord, application of the J60 ligand caused sustained increases in diaphragm output as measured through EMG in spontaneously breathing animals. This response was also verified using direct recordings of phrenic nerve discharge. Additionally, the DREADD ligand was able to produce an increase in inspiratory tidal volume in awake, freely behaving animals. These proof-of-concept studies provide a foundation for further development of this technology towards clinical application for restoring diaphragm activation in conditions such as cervical spinal cord injury.

### Targeted gene delivery to the phrenic motor pool

The intraspinal AAV delivery used here was based on previous studies demonstrating successful gene delivery to phrenic motoneurons^20^ ^24–26^. For example, mid-cervical spinal injections of an AAV5 vector encoding the lysosomal enzyme acid alpha-glucosidase (GAA) in animals with Pompe disease (*Gaa* null) effectively restores spinal GAA enzyme activity ^26^.

Spinal-delivered viral vectors have also been used to successfully drive local expression of channelrhodopsin-2 to enable light activation of diaphragm output^20^, and to drive expression of the astrocyte glutamate transporter GLT1 in the area of the phrenic motor nuclei^24^. Other methods that have been employed to drive gene expression in phrenic motoneurons include intrapleural- and intramuscular diaphragm injection of viral vectors^27^. Intrapleural delivery requires microinjection to the “pleural space” between the visceral pleura that lines the lungs and the parietal pleura that covers the thoracic cavity. This technique^28^ effectively targets phrenic motoneurons in rodent models of cervical spinal cord injury^29–31^ and Pompe disease^32^. Intramuscular diaphragm injection allows the vector to enter phrenic nerve terminals and reach phrenic motoneuron soma via retrograde movement^27^. Direct diaphragm injection allows for a relatively high target specificity, with the gene of interest expressed almost exclusively in phrenic motoneurons (although expression can also occur in diaphragm myofibers, depending on the promoter sequence used). In pilot experiments, we tested intrapleural and intramuscular diaphragm injections using AAV9 vectors encoding GFP (AAV9-CAG-GFP) or DREADD (AAV9-hSyn-HA-hM3D(Gq)-mCherry & AAV9-hSyn-DIO-hM3D(Gq)-mCherry). We did not, however, observe histological or physiological evidence of phrenic motoneuron transduction with these AAV9 vectors. Direct intraspinal injection^24,26^ was therefore used to introduce the hM3D(Gq) into the phrenic motor nucleus. While this enabled proof-of-concept for targeting DREADDs to the cervical spinal cord and phrenic motoneurons, the intrapleural or diaphragmatic injection delivery routes might ultimately prove better for selective phrenic motoneuron targeting. We predict that using different AAV serotypes or viruses with better retrograde movement (e.g., “AAV retro”) could optimize the targeting of phrenic motoneurons^33^.

### DREADD-mediated motoneuron activation

DREADD technology is widely used for studying brain and spinal cord neurons and networks^34,35^. Relatively few studies, however, have examined if and how DREADDs can be used to activate (or inhibit) lower motoneurons. Regarding the spinal cord, we are aware of only a few prior publications^36–39^. Two of these studies used pharmacologically selective actuator module (PSAM), a type of ionotropic chemogenetic receptor, to activate lumbar^36,37^ motoneurons, in mouse models of amyotrophic lateral sclerosis (ALS). In the remaining studies, excitatory DREADDs were applied to spinal motoneurons to improve axon regeneration following peripheral nerve injury^38,39^. A small but growing body of work has employed DREADDs to activate hypoglossal (XII) motoneurons in the brainstem^8^. Collectively, these studies show that once hM3D(Gq) is expressed in XII motoneurons, DREADD ligands will rapidly produce an increase in the EMG activation of tongue muscles^14,15^. This increase in tongue muscle output tends to manifest as an increase in the inspiratory-related activation and tonic discharge across the respiratory cycle. Since increased tongue muscle activation can promote patency of the upper airway, XII motoneuron DREADD expression has been suggested as a possible treatment for obstructive sleep apnea^12,14,16^. For the present study, the primary innovation is the first application of DREADD technology to phrenic motoneurons. This approach was highly effective at driving sustained activation of the diaphragm muscle, and the underlying mechanisms are discussed next.

### Chemogenetic stimulation of breathing

An important consideration is how DREADD-induced increases in the excitability of spinal neurons, including phrenic motoneurons, interacts with the endogenous neural control of breathing. Phrenic motoneurons receive a rhythmic, monosynaptic, glutamatergic synaptic input from medullary neurons. Acting via NMDA and AMPA receptors, this produces phrenic motoneuron depolarization and subsequent diaphragm muscle contraction^17^. Activating DREADDs on phrenic motoneurons should lower the threshold for activation via excitatory glutamatergic synaptic inputs, which would produce a greater output during the inspiratory phase. Alternatively, DREADD activation could directly lead to phrenic motoneuron action potentials even in the absence of synaptic input from the brainstem. This latter possibility could explain the tonic discharge (i.e., EMG output across the entire respiratory cycle) that was noted to occur after delivery of the DREADD ligand. Non-specific spinal cord DREADD expression, as occurred in the wild-type mice (e.g., **Figure 6**), would likely produce an increase in the excitability and/or activation of phrenic motoneurons as well as propriospinal neurons in the immediate vicinity. Neurophysiological^40^, as well as anatomical data^23,41^, confirm synaptic connections between mid-cervical interneurons and phrenic motoneurons, making it possible that DREADD activation of these interneurons impacted the diaphragm motor response in the wild-type mice.

The control of breathing is also impacted by well-established “closed loop” physiologic feedback mechanisms regulating lung volume and arterial blood gases^42,43^. For example, if DREADD-induced activation of the diaphragm leads to increased alveolar ventilation, and metabolic rate is not impacted, then arterial CO_2_ values will decrease and the overall neural drive to breathe will also decrease. Vagal afferent feedback corresponding to increased lung volume also has a powerful inhibitory impact on inspiration and therefore diaphragm activation. However, the sustained increase in diaphragm EMG and tidal volume that we observed following application of the DREADD ligand indicates that these mechanisms, if activated, were not sufficient to fully inhibit the increased phrenic motoneuron output. In this regard, our additional experiments in which direct recordings of bilateral phrenic nerve discharge are informative. These nerve recording experiments were done to enable direct evaluation of the impact of spinal DREADD activation on phrenic motor output while keeping arterial blood gases and lung volume constant. Under these more rigorously controlled conditions, intravenous delivery of the DREADD ligand produced a rapid and sustained increase in inspiratory burst amplitude in the phrenic nerve, and with no impact on the rate of the inspiratory bursts. The relative increase in inspiratory motor output was considerably greater in the phrenic nerve recording experiments (∼250% of baseline) as compared to the diaphragm EMG response in spontaneously breathing animals (∼100% of baseline). This may indicate that vagal and/or blood gas-related inhibitory mechanisms, as mentioned above, somewhat constrained the response to the DREADD ligand in the spontaneously breathing animal.

### Critique of methods

There are a few caveats that should be discussed. First, the precision of the AAV delivery could be improved by further refining spinal injection surgical techniques. In the current study, we used a stereotaxic frame and previously validated coordinates^26,44^ to guide the intraspinal AAV injections. However, we observed variability in the laterality (i.e., left vs. right side of the spinal cord) of mid-cervical mCherry expression as well as the physiological response to the DREADD ligand, particularly in the ChAT-Cre mice (e.g., **Figure 2**; **Table 1**). This could have occurred due to subtle variations of the positioning of the animal within the stereotaxic frame, and/or placement of the needle tip, leading to slight deviations for the desired coordinates between the left and right phrenic nuclei. Second, we did not unequivocally verify that the DREADD was expressed in phrenic motoneurons using retrograde labeling methods^28,45^. However, the phrenic motor nucleus has been well described in the mouse^46^ and the rat^47,48^, and the fluorophore (mCherry) expression observed in our experiments is very clearly in the expected location of phrenic motoneurons (**Figure 6b-c_i_; Figure S4**). Further, the robust increase in phrenic motor output after the DREADD ligand, particularly in the ChAT-Cre rat experiment (**Figure 5**) is further evidence of effective phrenic motoneuron targeting.

### Conclusion

Our data support the conclusion that cervical spinal cord directed chemogenetic methods can be used to produce sustained increases in phrenic motor output, diaphragm activation, and inspiratory tidal volume. Collectively, the data indicate that DREADDs should be directed exclusively to phrenic motoneurons vs. non-specific expression in the immediate vicinity. In this regard, improvement of the AAV delivery methods will increase the selectivity of the approach for more precise targeting of phrenic motoneurons. Concerning the “translational value” of this work, spinal cord chemogenetics may have application to clinical conditions associated with an inability to activate the diaphragm. For example, incomplete cervical spinal cord injury is a condition in which the bulbospinal synaptic inputs to phrenic motoneurons are interrupted. After incomplete cervical spinal cord injury, focal expression of an excitatory DREADD in phrenic motoneurons could be used to increase the excitability of these cells, thereby improving the efficacy of spared bulbospinal synaptic inputs which convey “inspiratory drive”.

## METHODS

### Animals

Experiments were carried out using C5/bl6, wild-type mice (Taconic), ChAT-Cre transgenic mice (B6.129S6-Chattm2(cre)Lowl/J; Jackson Laboratories), and ChAT-Cre transgenic rats (LE-Tg(Chat-Cre)5.1Deis; Rat Resource & Research Center). Animals were singly housed in a controlled environment (12 h light-dark cycle) with food and water *ad libitum*. All experiments were conducted in accordance with the NIH Guidelines Concerning the Care and Use of Laboratory Animals and were approved by the University of Florida Institutional Animal Care and Usage Committee (protocol #202107438). A full experimental timeline for mouse and rat experiments is shown in **Figure S6**, panels a and b, respectively.

### Adeno-associated viral vectors

All animals underwent intraspinal injections (see section below) of an AAV vector encoding the excitatory DREADD (hM3D(Gq)) under a human synapsin promoter. Wildtype mice received injections of AAV9-hSyn-HA-hM3D(Gq)-mCherry (titer: 2.44×10^13^ vg/mL) while ChAT-Cre mice and rats received injections of a similar construct with a double-floxed inverted open-reading frame (DIO) allowing for Cre-dependent transgene expression (AAV9-hSyn-DIO-hM3D(Gq)-mCherry; titer: 2.07×10^12^ vg/mL). The pAAV-hSyn-hM3D(Gq)-mCherry (Addgene plasmid # 50474; http://n2t.net/addgene:50474; RRID: Addgene_50474) and pAAV-hSyn-DIO-hM3D(Gq)-mCherry (Addgene plasmid # 44361; http://n2t.net/addgene:44361; RRID:Addgene_44361) transgene plasmids were gifts from the laboratory of Dr. Brian Roth at the University of North Carolina. Viral preparations were generated and titered by the University of Florida Powell Gene Therapy Center Vector Core Lab. Vectors were purified by iodixanol gradient centrifugation and anion-exchange chromatography as previously described^49^.

### Intraspinal injections

An adeno-associated viral vector (AAV) encoding the gene for the excitatory DREADD, hM3D(Gq) was delivered to the mid-cervical spinal cord. Mice were 6-10 weeks old (WT cohort: 7-9 weeks; ChAT-Cre cohort: 6-10 weeks) at the time of injection while ChAT-Cre rats were 2-5 months old. Surgery was performed under aseptic conditions. Mice were anesthetized with isoflurane (induction: 3-4% isoflurane; maintenance: 2-3% isoflurane in 100% O_2_) while rats were anesthetized with a mixture of ketamine (100 mg/kg) and xylazine (10 mg/kg) delivered intraperitoneally. Animals were placed prone on a circulating water heating pad to maintain body temperature. A longitudinal incision was made starting at the base of the skull and extending caudally. The underlying back musculature was opened from the base of the skull to spinal segment C6. Using a micro-curette, the muscle and connective tissue overlying laminae C3 to C5 were removed. A laminectomy of the C4 dorsal lamina exposed the dura mater below. A bilateral durotomy was then performed exposing the spinal cord. A Hamilton syringe (34-gauge needle) held in a Kopf stereotaxic frame was used to inject 1 µl of AAV9-hSyn-DIO-hM3D(Gq)-mCherry (ChAT-Cre mice and rats) or AAV9-hSyn-HA-hM3D(Gq)-mCherry (C57/bl6 mice), bilaterally into the ventral horns at C4. Injections were made 0.5 mm lateral to the spinal midline at a depth of 0.9 mm for mice^26^ and 1 mm lateral to midline at a depth of 1.5 mm for rats^44^. The needle was left to dwell for 5 minutes. Following injections, the overlying muscle and fascia were sutured with absorbable suture, the skin closed, and the animal returned to its home cage. Animals received a post-operative analgesia regiment of subcutaneous buprenorphine (1 mg/kg; slow-release formulation) and carprofen (5 mg/kg) for the first three days after surgery.

### Diaphragm EMG recordings

Recordings were conducted using wild-type (n = 11; n = 7 females) and ChAT-Cre mice (n = 9; n = 6 females; n = 1 excluded from analysis), 4-9 weeks following intraspinal injections of AAV-DREADD. Mice were anesthetized with 2-3% isoflurane in a closed chamber and then placed supine on a closed loop heating pad to maintain rectal temperature at 37 ± 0.5 °C (model 700 TC-1000, CWE Inc.). Mice spontaneously inhaled 2% isoflurane in 100% O_2_ for the duration of the experiment.

A laparotomy was performed and two sets of 50 µm tungsten wires were placed in the mid-costal region of the left and right hemidiaphragm. The tips of each wire were de-insulated, bent into small hooks, and inserted through the diaphragm approximately 3 mm apart. The recorded EMG signals were amplified (1000x) and filtered (100–1000 Hz) using a differential amplifier (A–M systems model 1700). Signals were digitized at 10 kS/s using a Power 1401 (CED, Cambridge, UK).

Once a stable plane of anesthesia was reached, mice underwent a 10-minute recording to establish baseline diaphragm EMG parameters. Subsequently, mice received injections of vehicle (100 µl of saline delivered intraperitoneally (IP)) followed by a 20-minute recording. Mice then received an intraperitoneal injection of the selective DREADD agonist, JHU37160 (J60; 0.1 mg/kg, HB6261, HelloBio), and recordings continued for 90 minutes. At the conclusion of each experiment, mice underwent transcardial perfusion with saline followed by 4% paraformaldehyde. Following perfusion, spinal cords were harvested for histological analysis.

### J60 control experiments

A small cohort of animals (n = 2 C57/bl mice; n = 3 Sprague Dawley rats) was used to assess the impact of J60 (0.1 mg/kg) on diaphragm EMG activity in the absence of hM3D(Gq) expression. The animals used in this study include n = 2 C57/bl mice that had undergone intrapleural injection (i.e., injection to the thoracic cavity) of an AAV9 construct encoding the red fluorescent protein, mCherry and n = 3 vector naïve Sprague Dawley rats.

Recordings in mice proceeded as described above (see *Diaphragm EMG recordings*). In rat recordings, rats were induced with 3% isoflurane in 100% O_2_ and moved onto a closed-loop heating pad set to maintain rectal temperature at 37 ± 1°C (model 700 TC-1000, CWE Inc.). Rats were tracheotomized and ventilated (Model 683; Harvard Apparatus Inc.) with a gas mixture of 50% O_2_, and 1% CO_2_, balanced with N_2_. End-tidal CO_2_ was maintained at 45-47 mmHg throughout the experimental protocol (Capnogard; Novametrix). Rats were converted from isoflurane to urethane anesthesia (2.1 g/kg at 6 mL/hr; IV). At the competition of urethane dosing lactated Ringer’s was administered (2 mL/h; IV) to keep the animal hydrated and ensure the catheter remained viable for J60 administration. A femoral artery catheter (polyethylene tubing; PE 50; Intramedic) was placed to enable monitoring of arterial blood pressure via a transducer amplifier (TA-100, CWE).

At the beginning of the experimental period, rats underwent a 10-minute recording to establish baseline diaphragm EMG parameters. This was followed by an IV injection of vehicle (0.6 mL of saline) and a subsequent 20-minute recording. Next, rats received an IV infusion of the J60 agonist (0.1 mg/kg), and the recording continued for 90 minutes. At the end of the experiment, rats were euthanized via an overdose of pentobarbital sodium and phenytoin sodium (150 mg/kg) given intravenously. Death was confirmed by thoracotomy once breathing had ceased, and a heartbeat was no longer detectable.

### Whole body plethysmography

ChAT-Cre rats (n = 9; n = 3 females) were studied using flow-through whole-body plethysmography 14-16 weeks after intraspinal delivery of AAV9-hSyn-DIO-hM3D(Gq)-mCherry, as described above. A tail vein catheter was placed to allow for intravenous infusion (IV) of the J60 ligand and vehicle. An IV catheter was externalized via a port in the plethysmograph allowing for IV infusion during recording without handling the animal or opening the plethysmograph. Unanesthetized rats were sealed into the Plexiglas plethysmograph with airflow maintained at 6 L/min for the duration of the recording. The recording protocol consisted of a 40-minute acclimation period (inspired air: 21% O_2_, 79% N_2_), followed by a 7-minute ventilatory challenge (10% O_2_, 7% CO_2_, 83% N_2_) and a 10-minute normoxic recovery period (21% O_2_, 79% N_2_). Subsequently, rats underwent a 20-minute long, pre-vehicle, baseline under normoxic conditions (21% O_2_, 79% N_2_) followed by a 2-minute-long intravenous infusion of the J60 vehicle (saline; 0.6 mL). Following vehicle infusion recording continued for 30 minutes followed by a 7-minute ventilatory challenge (10% O_2_, 7% CO_2_, 83% N_2_) and 10 minutes of normoxic breathing (21% O_2_, 79% N_2_). After an additional 20-minute pre-J60 baseline (21% O_2_, 79% N_2_), an intravenous infusion of the J60 ligand was given (0.1 mg/ml dose; 2 minutes long; final volume standardized to 0.6 mL) and recordings continued for 30-minute followed by a final ventilatory challenge (10% O_2_, 7% CO_2_, 83% N_2_). The ventilatory challenges were performed to assess the ability to increase breathing.

### Phrenic nerve recordings

Two-to-eight-weeks following plethysmography recordings, bilateral phrenic nerve recordings were performed. This procedure was done to directly assess the effect of DREADD activation on phrenic motor output under rigorously controlled experimental conditions. Anesthesia was induced by placing the rat in a closed chamber to inhale 3% isoflurane in 100% O_2_. Rats were then moved onto a closed-loop heating pad set to maintain rectal temperature at 37 ± 1°C (model 700 TC-1000, CWE Inc.). Isoflurane anesthesia was maintained using a nose cone. Once a surgical plane of anesthesia was reached as evidenced by loss of corneal reflexes and hindlimb withdrawal, rats were tracheotomized and ventilated (VentElite, model 55-7040; Harvard Apparatus Inc.) with a gas mixture of 50% O_2_, 1% CO_2_, balanced with N_2_. End-tidal CO_2_ was maintained at 45-47 mmHg throughout the surgery and experimental protocol (Capnogard; Novametrix). Ventilator frequency was maintained between 65 and 75 breaths/min, and tidal volume was set at 7 mL/kg^50^. The vagus nerves were transected bilaterally to prevent entrainment of phrenic efferent output with the ventilator.

A tail vein catheter was placed to allow for intravenous infusion of urethane anesthesia, supplementary fluids, and the J60 ligand. Rats were slowly converted from inhaled isoflurane to urethane anesthesia (2.1 g/kg at 6 mL/hr; IV). During this conversion, the depth of anesthesia was consistently monitored by evaluating the pedal withdrawal reflex. Following administration of the full urethane dose, a mixture of 8.4% sodium bicarbonate and lactated Ringer’s was administered (2 mL/h; IV) to maintain acid-base balance. To prevent movements and EMG contamination of the phrenic neurogram pancuronium bromide was administered (3 mg/kg IV, Sigma-Aldrich, St Louis) to achieve neuromuscular blockade. A catheter (polyethylene tubing; PE 50; Intramedic) was placed in the femoral artery to enable monitoring of arterial blood pressure via a transducer amplifier (TA-100, CWE) and allow withdrawal of arterial blood samples (65 μL) for measurement of partial pressure of CO_2_ (PaCO_2_) and O_2_ (PaO_2_), pH, and base excess (ABL 90 Flex, Radiometer; Copenhagen, Denmark).

The phrenic nerves were exposed bilaterally using a dorsal approach as described previously^51,52^. Briefly, a midline incision was made at the base of the skull extending to spinal level T2. The muscles connecting the shoulder blades to the spinal column were separated to expose the phrenic nerves. The phrenic nerves were isolated, cut distal to the spinal cord, and suctioned into custom-made glass electrodes filled with 0.9% saline solution. Phrenic nerve activity was amplified (10 kHz) using a differential AC amplifier (Model 1700, A-M systems, Everett, WA), band-pass filtered (100Hz-3 kHz), and digitized at 25ks/second (Power 1401, CED).

At the beginning of the experiment, the apneic threshold was determined by slowly reducing the inspired CO_2_ until phrenic nerve inspiratory activity ceased for 60 seconds. The recruitment threshold was established by slowly increasing the inspired CO_2_ until phrenic bursting returned. The end-tidal CO_2_ (ETCO_2_) was then maintained 2-3 mmHg above the recruitment threshold for the duration of the experiment. After achieving a stable phrenic nerve recording and blood gases a 15-minute-long baseline recording was collected (50% O_2_, 3% CO_2_) followed by a brief, 5-minute exposure to hypoxia (11.5% O_2_, 3% CO_2_) and 10–15-minute recovery period (50% O_2_, 3% CO_2_). Subsequently, intravenous infusion of vehicle (saline) was given followed by a 15-minute recording period. The J60 ligand (0.1 mg/kg) was then administered intravenously over a 2-minute infusion period followed by a 100-minute recording period.

Arterial blood samples were collected at specific intervals: initially at baseline, during the last minute of each hypoxia episode, 15 minutes post vehicle administration, and subsequently at 20-, 40,-60-, 80-, and 100-minutes post J60 administration. Baseline blood gas values served as references to assess if further arterial samples were isocapnic. To keep end-tidal CO_2_ and PaCO_2_ near baseline (within ± 2.0 mmHg), minor adjustments to inspired CO_2_ and ventilation rate were made as needed. PaO_2_ was kept above 150 mmHg, except during hypoxia; if it dropped below, O_2_ intake was increased by 5%, and a new blood sample was analyzed within 5 minutes.

At the end of the experiment, rats were exposed to a second 5-min episode of hypoxia (11.5% O_2_) followed by a brief “maximal” chemoreceptor challenge induced by switching off the mechanical ventilator until the animal exhibited a “gasping-like” phrenic discharge pattern (approximately 20-30 seconds). If the increase in phrenic nerve amplitude in response to the “maximal” challenge was lower than the response observed during either hypoxic episode, it was considered a sign of deteriorating nerve-electrode contact, and the preparation was excluded from all formal analyses. Rats were then perfused transcardially with heparinized saline followed by 4% paraformaldehyde and spinal cords were harvested for histological analysis.

### Histology

Spinal cords were harvested and placed in 4% paraformaldehyde for 24 hours. The cords were subsequently moved to a cryo-protecting solution (30% sucrose in 1x PBS) for a minimum of three days. Cervical and thoracic spinal cords were blocked in optimal cutting temperature media and cryosectioned at 20 µm. The viral constructs included a red fluorescent protein (mCherry) fused to the hM3D(Gq) DREADD which allowed evaluation of DREADD expression by assessing mCherry expression via fluorescence microscopy.

We performed a qualitative assessment of mCherry expression in the mid-cervical spinal cord. One intact section from the middle of each spinal segment (C3-C6) was chosen as a representative section and underwent assessment. Sections were segmented into the following quadrants: left dorsal, right dorsal, left ventral, and right ventral. The quadrant was scored as “positive” if mCherry positive neurons or fibers were observed; otherwise, the sub-segment was marked “negative” (see **Figure S4** for example). The entirety of the grey matter from each section was analyzed for all animals, whether wild-type or ChAT-Cre. Although ChAT-Cre expression was expected to be limited primarily to motoneurons, which are the predominant ChAT-positive neuronal subtype in the spinal cord, there is also evidence of ChAT-positive interneuron populations^53–55^ which we also wished to capture in our analysis. Results were compiled into a summary table showing the total positive counts by animal cohort, spinal segment, and quadrant (see Results section; **Table 1**). Animals that showed no positive mCherry labeling in the C3-C6 cord were excluded from analysis.

### Data analysis

Custom MATLAB (MathWorks; Natick, MA) scripts were created, and are available upon request. These scripts were used to analyze diaphragm EMG, phrenic nerve, and plethysmography waveforms. EMG signals were digitally filtered using a second-order, bandpass Butterworth filter (100– 1000 Hz) and then rectified and integrated by taking the absolute value of the signal followed by applying a moving median filter (50 ms time constant for mice; 75 ms time constant for rats) and moving average filter (50 ms time constant for mice; 175 ms time constant for rats). The script identified each EMG burst and calculated peak amplitude, minimum amplitude (tonic activity), and AUC for each burst which was then averaged across animals and compared across experimental conditions.

Phrenic nerve signals were digitally filtered using a second-order, bandpass Butterworth filter (100–3 kHz) and then rectified and integrated by taking the absolute value of the signal followed by applying a moving median filter (50 ms time constant) and moving average filter (50 ms time constant). The analysis script calculated the peak phrenic burst amplitude and minimum amplitude for each burst which was then averaged across animals and compared across experimental conditions. Systolic (SP), diastolic (DP), and mean arterial blood pressure (MAP; formula: MAP = DP + 1/3 (SP - DP)) along with instantaneous heart rate were calculated from the arterial pressure trace.

In plethysmography experiments, airflow pressure, chamber temperature, chamber humidity, barometric pressure, and animal body temperature were used to calculate respiratory frequency, tidal volume, and ventilation via a custom MATLAB script. Tidal volume was calculated using the Drorbaugh and Fenn equation^56^.

Statistical analyses were performed using SigmaPlot 14 (Systat Software) and R (The R Foundation for Statistical Computing; version 4.3.1). In mouse diaphragm EMG studies, one-way repeated measure analysis of variance (ANOVA) was used to statistically compare diaphragm EMG peak amplitude, area under the curve, tonic activity, and heart rate across time before and after J60 application. Paired t-tests were used to compare left and right hemidiaphragm EMG peak amplitude, area under the curve, tonic activity, and heart rate between ChAT-Cre and wild-type mice at the 30-minute post-J60 administration time point. Differences in mortality between wild-type and ChAT-Cre mice post-J60 application were assessed using Pearson’s Chi-squared test with Yates’ continuity correction using the chisq.test function in R. In instances where animals did not survive the entire duration of the anesthetized recording, data up until the time point preceding their death was included. In control EMG experiments, one-way RM ANOVA was used to compare EMG peak responses across baseline, sham injection, and J60 administration. These data were also assessed normalized to baseline, in which case EMG peak responses after sham injection and J60 application were compared using paired t-tests. In plethysmography experiments, two-way repeated measures ANOVA was used to compare raw and normalized tidal volume, respiratory frequency, and minute ventilation across time and treatment (saline vs J60). Paired t-tests were used to compare responses to hypercapnic-hypoxic ventilatory challenges across treatments. One-way RM ANOVA was used to compare phrenic peak amplitude, systolic and diastolic blood pressures, mean arterial blood pressure, and respiratory rate across time for phrenic nerve recordings. The relationship between time post-AAV injection and average phrenic response to J60 was assessed for ChAT-Cre rats using the cor.test function in R to run a Pearson’s product moment correlation. Both male and female animals were included in this study to improve the generalizability of the results. However, we were not adequately powered for sex comparisons and therefore did not perform any statistical analysis to assess sex differences.

In cases of significant main effects, the Tukey post-hoc test was used to assess differences between individual time points. For instances where data failed to meet general linear model assumptions (i.e., normality, homogeneity of variances), nonparametric equivalents of the previously mentioned statistical tests were used. Data were considered statistically significant when p ≤ 0.05. The mean data are presented along with the standard error of the mean.

## Abbreviations

DREADD: (designer receptors exclusively activated by designer drugs)
EMG: (electromyography)
AUC: (area under the curve)
J60: (JHU37160)
AAV: (adeno-associated virus)
VT: (tidal volume)
ChAT: (choline acetyltransferase)
Cre: (Cre recombinase)

## Notes

### Competing Interest Statement

The authors have declared no competing interest.

### Summary of Updates

Minor text edits. Additional control experiments added.

## REFERENCES

1 Berlowitz, D. J., Wadsworth, B. & Ross, J. Respiratory problems and management in people with spinal cord injury. Breathe (Sheff) 12, 328–340 (2016). 10.1183/20734735.012616

2 Brown, R., DiMarco, A. F., Hoit, J. D. & Garshick, E. Respiratory dysfunction and management in spinal cord injury. Respir Care 51, 853–868;discussion 869-870 (2006).

3 Perrin, C., Unterborn, J. N., Ambrosio, C. D. & Hill, N. S. Pulmonary complications of chronic neuromuscular diseases and their management. Muscle Nerve 29, 5–27 (2004). 10.1002/mus.10487

4 Mehta, S. Neuromuscular disease causing acute respiratory failure. Respir Care 51, 1016–1021; discussion 1021-1013 (2006).

5 Burakgazi, A. Z. & Höke, A. Respiratory muscle weakness in peripheral neuropathies. J Peripher Nerv Syst 15, 307–313 (2010). 10.1111/j.1529-8027.2010.00293.x

6 Fuller, D. D. et al. The respiratory neuromuscular system in Pompe disease. Respir Physiol Neurobiol 189, 241–249 (2013). 10.1016/j.resp.2013.06.007

7 Jordan, A. S. & White, D. P. Pharyngeal motor control and the pathogenesis of obstructive sleep apnea. Respir Physiol Neurobiol 160, 1–7 (2008). 10.1016/j.resp.2007.07.009

8 Doyle, B. M. et al. Gene delivery to the hypoglossal motor system: preclinical studies and translational potential. Gene Ther (2021). 10.1038/s41434-021-00225-1

9 Zhu, H. & Roth, B. L. DREADD: a chemogenetic GPCR signaling platform. Int J Neuropsychopharmacol 18 (2014). 10.1093/ijnp/pyu007

10 Alexander, G. M. et al. Remote control of neuronal activity in transgenic mice expressing evolved G protein-coupled receptors. Neuron 63, 27–39 (2009). 10.1016/j.neuron.2009.06.014

11 Armbruster, B. N., Li, X., Pausch, M. H., Herlitze, S. & Roth, B. L. Evolving the lock to fit the key to create a family of G protein-coupled receptors potently activated by an inert ligand. Proc Natl Acad Sci U S A 104, 5163–5168 (2007). 10.1073/pnas.0700293104

12 Fleury Curado, T., et al. Chemogenetic stimulation of the hypoglossal neurons improves upper airway patency. Scientific reports 7, 44392 (2017). 10.1038/srep44392

13 Fleury Curado, T. A., et al. Silencing of Hypoglossal Motoneurons Leads to Sleep Disordered Breathing in Lean Mice. Front Neurol 9, 962 (2018). 10.3389/fneur.2018.00962

14 Fleury Curado, T., et al. Designer Receptors Exclusively Activated by Designer Drugs Approach to Treatment of Sleep-disordered Breathing. American journal of respiratory and critical care medicine 203, 102–110 (2021). 10.1164/rccm.202002-0321OC

15 Singer, M. L. et al. Chemogenetic activation of hypoglossal motoneurons in a mouse model of Pompe disease. Journal of neurophysiology 128, 1133–1142 (2022). 10.1152/jn.00026.2022

16 Horton, G. A. et al. Activation of the Hypoglossal to Tongue Musculature Motor Pathway by Remote Control. Sci Rep 7, 45860 (2017). 10.1038/srep45860

17 Fuller, D. D., Rana, S., Smuder, A. J. & Dale, E. A. The phrenic neuromuscular system. Handbook of clinical neurology 188, 393–408 (2022). 10.1016/B978-0-323-91534-2.00012-6

18 DiMarco, A. F. & Kowalski, K. E. Activation of inspiratory muscles via spinal cord stimulation. Respiratory physiology & neurobiology 189, 438–449 (2013). 10.1016/j.resp.2013.06.001

19 Jensen, V. N., Seedle, K., Turner, S. M., Lorenz, J. N. & Crone, S. A. V2a Neurons Constrain Extradiaphragmatic Respiratory Muscle Activity at Rest. eNeuro 6 (2019). 10.1523/ENEURO.0492-18.2019

20 Alilain, W. J. et al. Light-induced rescue of breathing after spinal cord injury. The Journal of neuroscience : the official journal of the Society for Neuroscience 28, 11862–11870 (2008). 10.1523/JNEUROSCI.3378-08.2008

21 Satkunendrarajah, K., Karadimas, S. K., Laliberte, A. M., Montandon, G. & Fehlings, M. G. Cervical excitatory neurons sustain breathing after spinal cord injury. Nature 562, 419–422 (2018). 10.1038/s41586-018-0595-z

22 Jensen, V. N., Alilain, W. J. & Crone, S. A. Role of Propriospinal Neurons in Control of Respiratory Muscles and Recovery of Breathing Following Injury. Front Syst Neurosci 13, 84 (2019). 10.3389/fnsys.2019.00084

23 Lane, M. A. Spinal respiratory motoneurons and interneurons. Respiratory physiology & neurobiology 179, 3–13 (2011). 10.1016/j.resp.2011.07.004

24 Li, K. et al. Overexpression of the astrocyte glutamate transporter GLT1 exacerbates phrenic motor neuron degeneration, diaphragm compromise, and forelimb motor dysfunction following cervical contusion spinal cord injury. J Neurosci 34, 7622–7638 (2014). 10.1523/jneurosci.4690-13.2014

25 Li, K. et al. GLT1 overexpression in SOD1(G93A) mouse cervical spinal cord does not preserve diaphragm function or extend disease. Neurobiology of disease 78, 12–23 (2015). 10.1016/j.nbd.2015.03.010

26 Qiu, K., Falk, D. J., Reier, P. J., Byrne, B. J. & Fuller, D. D. Spinal delivery of AAV vector restores enzyme activity and increases ventilation in Pompe mice. Molecular therapy : the journal of the American Society of Gene Therapy 20, 21–27 (2012). 10.1038/mt.2011.214

27 Thakre, P. P., Rana, S., Benevides, E. S. & Fuller, D. D. Targeting drug or gene delivery to the phrenic motoneuron pool. Journal of neurophysiology 129, 144–158 (2023). 10.1152/jn.00432.2022

28 Mantilla, C. B., Zhan, W. Z. & Sieck, G. C. Retrograde labeling of phrenic motoneurons by intrapleural injection. J Neurosci Methods 182, 244–249 (2009). 10.1016/j.jneumeth.2009.06.016

29 Gransee, H. M., Zhan, W. Z., Sieck, G. C. & Mantilla, C. B. Targeted delivery of TrkB receptor to phrenic motoneurons enhances functional recovery of rhythmic phrenic activity after cervical spinal hemisection. PLoS One 8, e64755 (2013). 10.1371/journal.pone.0064755

30 Martínez-Gálvez, G. et al. TrkB gene therapy by adeno-associated virus enhances recovery after cervical spinal cord injury. Exp Neurol 276, 31–40 (2016). 10.1016/j.expneurol.2015.11.007

31 Gransee, H. M., Gonzalez Porras, M. A., Zhan, W. Z., Sieck, G. C. & Mantilla, C. B. Motoneuron glutamatergic receptor expression following recovery from cervical spinal hemisection. J Comp Neurol 525, 1192–1205 (2017). 10.1002/cne.24125

32 Keeler, A. M. et al. Intralingual and Intrapleural AAV Gene Therapy Prolongs Survival in a SOD1 ALS Mouse Model. Mol Ther Methods Clin Dev 17, 246–257 (2020). 10.1016/j.omtm.2019.12.007

33 Tervo, D. G. et al. A Designer AAV Variant Permits Efficient Retrograde Access to Projection Neurons. Neuron 92, 372–382 (2016). 10.1016/j.neuron.2016.09.021

34 Smith, K. S., Bucci, D. J., Luikart, B. W. & Mahler, S. V. Dreadds: Use and application in behavioral neuroscience. Behav Neurosci 135, 89–107 (2021). 10.1037/bne0000433

35 Roth, B. L. DREADDs for Neuroscientists. Neuron 89, 683–694 (2016). 10.1016/j.neuron.2016.01.040

36 Ouali Alami, N., et al. Multiplexed chemogenetics in astrocytes and motoneurons restore blood-spinal cord barrier in ALS. Life Sci Alliance 3 (2020). 10.26508/lsa.201900571

37 Saxena, S. et al. Neuroprotection through excitability and mTOR required in ALS motoneurons to delay disease and extend survival. Neuron 80, 80–96 (2013). 10.1016/j.neuron.2013.07.027

38 Jaiswal, P. B., Mistretta, O. C., Ward, P. J. & English, A. W. Chemogenetic Enhancement of Axon Regeneration Following Peripheral Nerve Injury in the SLICK-A Mouse. Brain Sci 8 (2018). 10.3390/brainsci8050093

39 Jaiswal, P. B. & English, A. W. Chemogenetic enhancement of functional recovery after a sciatic nerve injury. Eur J Neurosci 45, 1252–1257 (2017). 10.1111/ejn.13550

40 Streeter, K. A. et al. Mid-cervical interneuron networks following high cervical spinal cord injury. Respir Physiol Neurobiol 271, 103305 (2020). 10.1016/j.resp.2019.103305

41 Lane, M. A. et al. Cervical prephrenic interneurons in the normal and lesioned spinal cord of the adult rat. The Journal of comparative neurology 511, 692–709 (2008). 10.1002/cne.21864

42 Molkov, Y. I., Rubin, J. E., Rybak, I. A. & Smith, J. C. Computational models of the neural control of breathing. Wiley Interdiscip Rev Syst Biol Med 9 (2017). 10.1002/wsbm.1371

43 Dempsey, J. A. & Welch, J. F. Control of Breathing. Semin Respir Crit Care Med 44, 627–649 (2023). 10.1055/s-0043-1770342

44 McGuire, M., Zhang, Y., White, D. P. & Ling, L. Phrenic long-term facilitation requires NMDA receptors in the phrenic motonucleus in rats. J Physiol 567, 599–611 (2005). 10.1113/jphysiol.2005.087650

45 Rana, S., Zhan, W. Z., Sieck, G. C. & Mantilla, C. B. Cervical spinal hemisection alters phrenic motor neuron glutamatergic mRNA receptor expression. Experimental neurology 353, 114030 (2022). 10.1016/j.expneurol.2022.114030

46 Qiu, K., Lane, M. A., Lee, K. Z., Reier, P. J. & Fuller, D. D. The phrenic motor nucleus in the adult mouse. Experimental neurology 226, 254–258 (2010). 10.1016/j.expneurol.2010.08.026

47 Rana, S., Sieck, G. C. & Mantilla, C. B. Heterogeneous glutamatergic receptor mRNA expression across phrenic motor neurons in rats. Journal of neurochemistry 153, 586–598 (2020). 10.1111/jnc.14881

48 Rana, S., Mantilla, C. B. & Sieck, G. C. Glutamatergic input varies with phrenic motor neuron size. J Neurophysiol 122, 1518–1529 (2019). 10.1152/jn.00430.2019

49 Zolotukhin, S. et al. Production and purification of serotype 1, 2, and 5 recombinant adeno-associated viral vectors. Methods 28, 158–167 (2002). 10.1016/s1046-2023(02)00220-7

50 Lee, K. Z., Sandhu, M. S., Dougherty, B. J., Reier, P. J. & Fuller, D. D. Hypoxia triggers short term potentiation of phrenic motoneuron discharge after chronic cervical spinal cord injury. Exp Neurol 263, 314–324 (2015). 10.1016/j.expneurol.2014.10.002

51 Thakre, P. P. & Fuller, D. D. Pattern sensitivity of ampakine-hypoxia interactions for evoking phrenic motor facilitation in anesthetized rat. J Neurophysiol 131, 216–224 (2024). 10.1152/jn.00315.2023

52 Thakre, P. P., Sunshine, M. D. & Fuller, D. D. Ampakine pretreatment enables a single hypoxic episode to produce phrenic motor facilitation with no added benefit of additional episodes. J Neurophysiol 126, 1420–1429 (2021). 10.1152/jn.00307.2021

53 Gotts, J., Atkinson, L., Yanagawa, Y., Deuchars, J. & Deuchars, S. A. Co-expression of GAD67 and choline acetyltransferase in neurons in the mouse spinal cord: A focus on lamina X. Brain Res 1646, 570–579 (2016). 10.1016/j.brainres.2016.07.001

54 Alkaslasi, M. R. et al. Single nucleus RNA-sequencing defines unexpected diversity of cholinergic neuron types in the adult mouse spinal cord. Nat Commun 12, 2471 (2021). 10.1038/s41467-021-22691-2

55 Mesnage, B. et al. Morphological and functional characterization of cholinergic interneurons in the dorsal horn of the mouse spinal cord. J Comp Neurol 519, 3139–3158 (2011). 10.1002/cne.22668

56 Drorbaug, J. E. & Fenn, W. O. A BAROMETRIC METHOD FOR MEASURING VENTILATION IN NEWBORN INFANTS. Pediatrics 16, 81–87 (1955). 10.1542/peds.16.1.81

