## Supplemental files for "Chemogenetic stimulation of phrenic motor output and diaphragm activity"

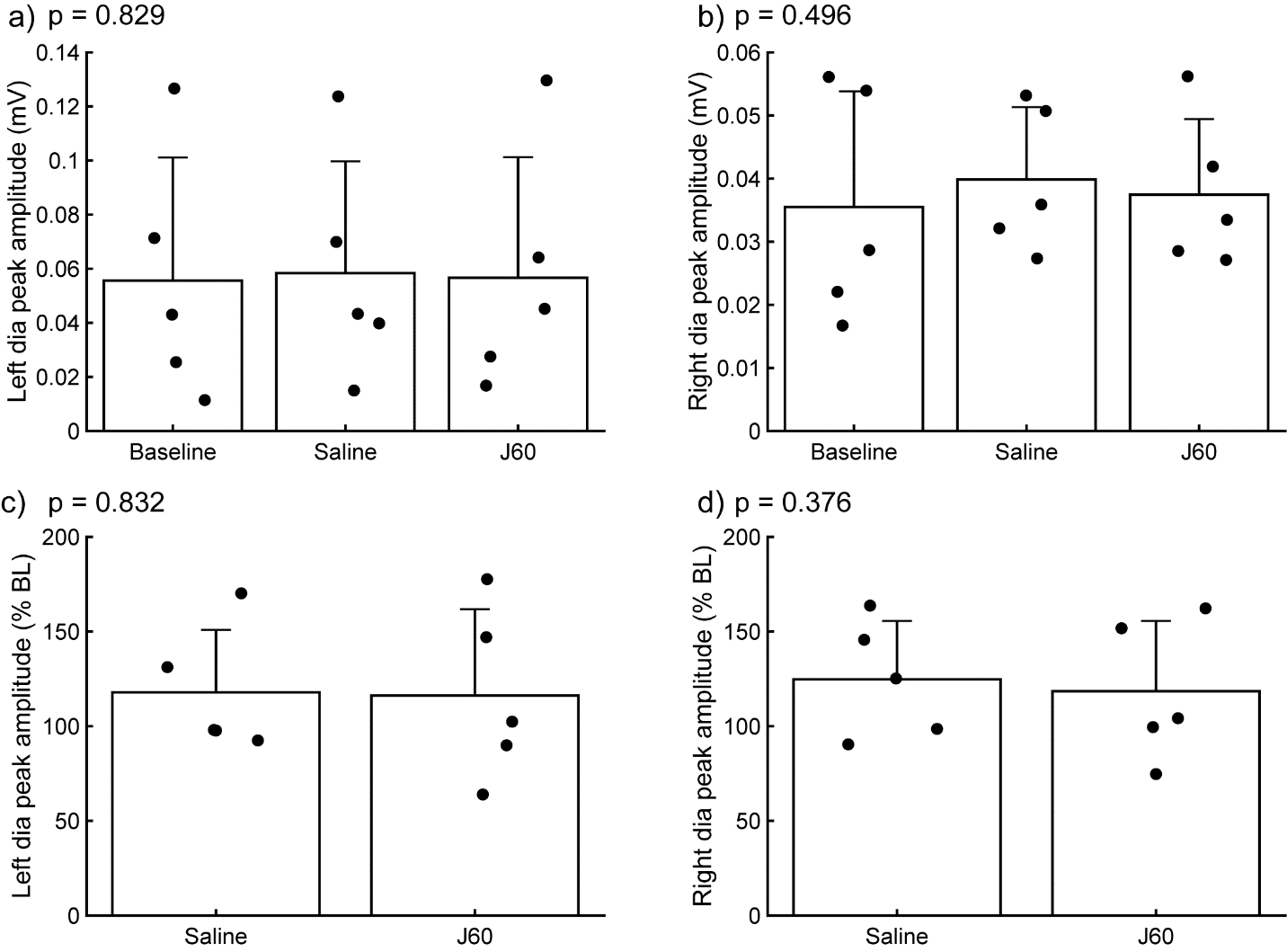

**Supplemental Figure 1 (S1). *Impact of J60 application on diaphragm EMG in the absence of hM3D(Gq) expression.*** Summary plots of a combined mouse (n = 2) and rat (n = 3) data showing the impact of J60 application (0.1 mg/kg) on diaphragm EMG peak amplitude in animals not expressing the hM3D(Gq) DREADD. Mean diaphragm EMG responses during baseline, saline (sham injection), and following J60 administration for the left hemidiaphragm are shown in panels a and b, respectively. Mean values normalized to baseline are shown in panels c (left hemidiaphragm) and d (right hemidiaphragm). Raw values (a-b) were assessed by one-way RM ANOVA while baseline normalized values (c-d) were assessed via paired t-tests. No statistically significant differences were detected in either hemidiaphragm across experimental periods. P-values are displayed next to each panel legend. Error bars depict ± 1 SEM.

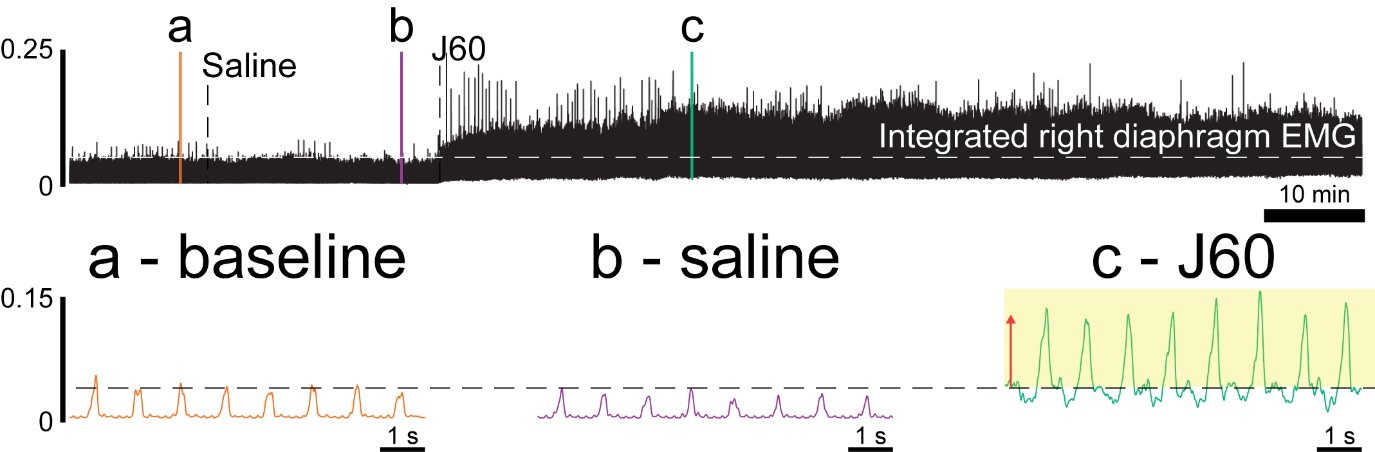

**Supplemental Figure 2 (S2). *Impact of DREADD activation on diaphragm EMG in ChAT-Cre rats.*** Example trace of rectified and integrated diaphragm EMG activity from a ChAT-Cre rat that had previously undergone injections of AAV9-hSyn-DIO-hM3D(Gq)-mCherry into the C4 ventral horns before and after application of the DREADD ligand, J60. Callout panels below show example activity at baseline (a), after injection of vehicle (b), and after application of the DREADD ligand, J60 (c).

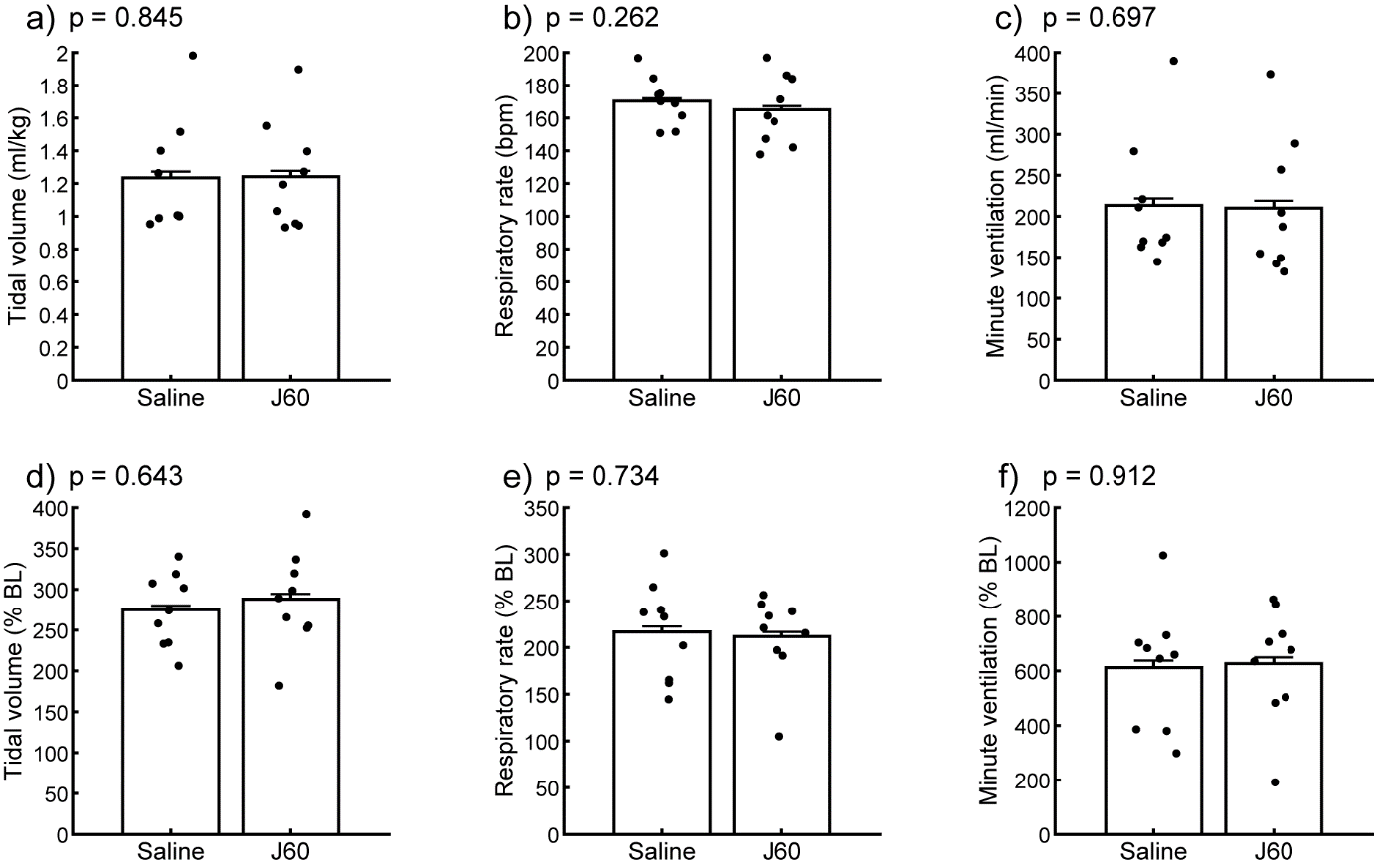

**Supplemental Figure 3 (S3). *Ventilatory responses to hypercapnic-hypoxic respiratory challenge*.** Summary plots (n = 9; n = 3 females) showing the impact of the J60 DREADD ligand on tidal volume, respiratory rate, and minute ventilation during a hypercapnic-hypoxic ventilatory challenge (panels a-c). The normalized values (% of baseline) are shown in panels d-f. No difference was detected in any of the ventilatory parameters across the sham vs J60 condition during the hypercapnic-hypoxic ventilatory challenge. Paired t-tests were performed on the data in each panel. P-values are displayed next to each panel legend. Error bars depict ± 1 SEM.

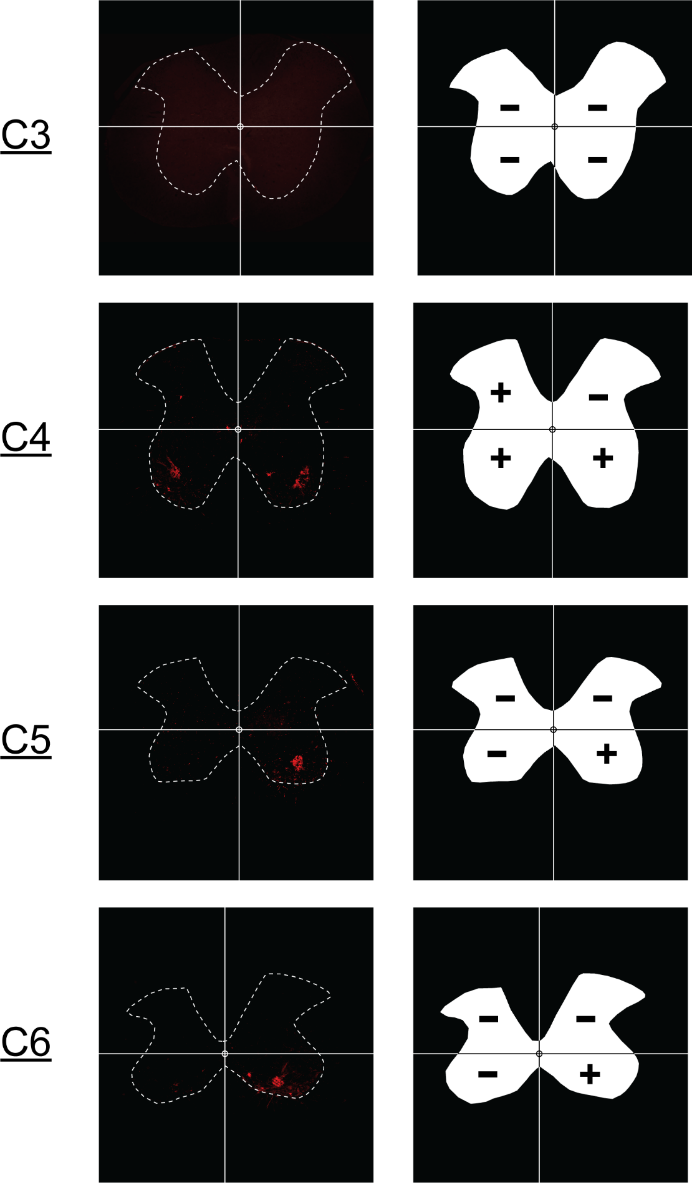

**Supplemental Figure 4 (S4). *Example quantification of mCherry expression in the C3-C6 spinal cord.*** Mid-cervical spinal cord images from ChAT-Cre animal (left) with positive mCherry expression. A grid has been placed on each image to highlight the quadrant system (i.e., left dorsal, right dorsal, left ventral, right ventral) used to score the tissue. The right column shows an outline of each section on the left with a score for each quadrant based on mCherry expression. A given quadrant was scored with a “+” if it contained any mCherry positive neurons or fibers else it was given a “-“ signifying no mCherry expression.

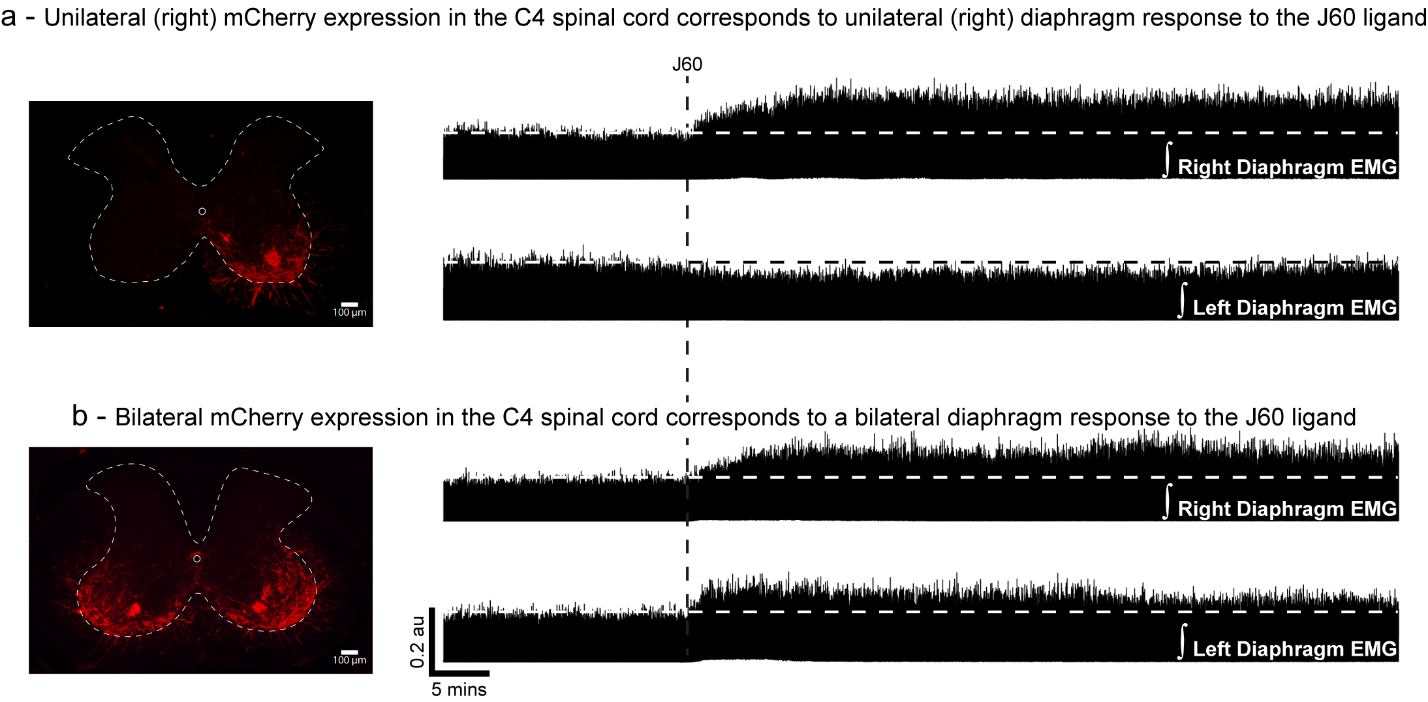

**Supplemental Figure 5 (S5). *mCherry expression in the mid-cervical ventral horn corresponds with the laterality of the diaphragm EMG DREADD response*.** Example spinal histology (left) from two ChAT-Cre mice showing unilateral (a) and bilateral (b) expression of mCherry (red) in the ventral horn(s) and corresponding diaphragm EMG traces. Unilateral ventral horn mCherry expression (a) results in DREADD responses that were limited to the ipsilateral hemidiaphragm. Animals with bilateral ventral horn DREADD expression responded to DREADD activation with a bilateral increase in diaphragm output.

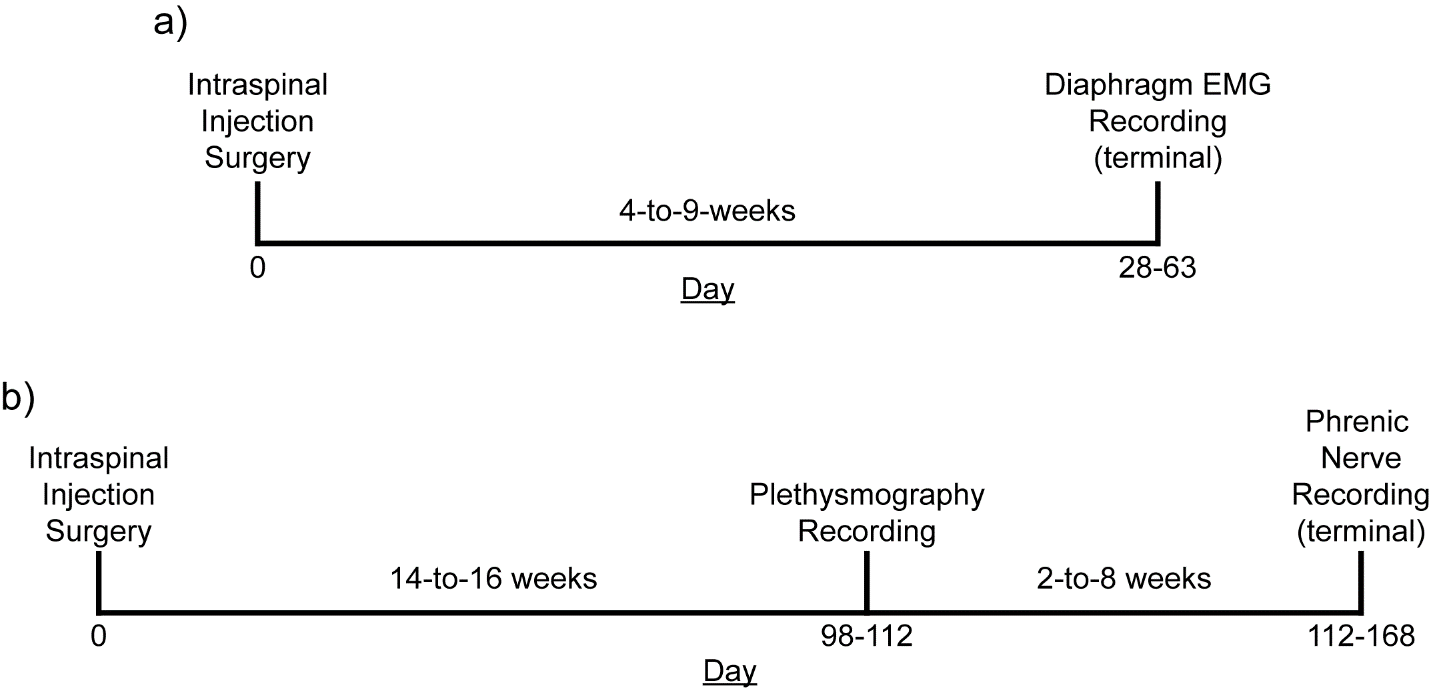

**Supplemental Figure 6 (S6). *Experimental timelines.*** Timelines of mouse (a) and rat (b) studies. Both cohorts of animals underwent an initial surgery to introduce an AAV vector encoding the excitatory DREADD, hM3D(Gq) in the ventral mid-cervical spinal cord bilaterally. Mice incubated for 4-to-9 weeks before undergoing terminal diaphragm EMG recordings. Rats incubated for 14-to-16 weeks before undergoing plethysmography recordings, 2-to-8 weeks later rats underwent terminal phrenic nerve recordings. In all experiments, baseline parameters were established followed by an infusion of vehicle, and subsequently the selective DREADD ligand, J60.

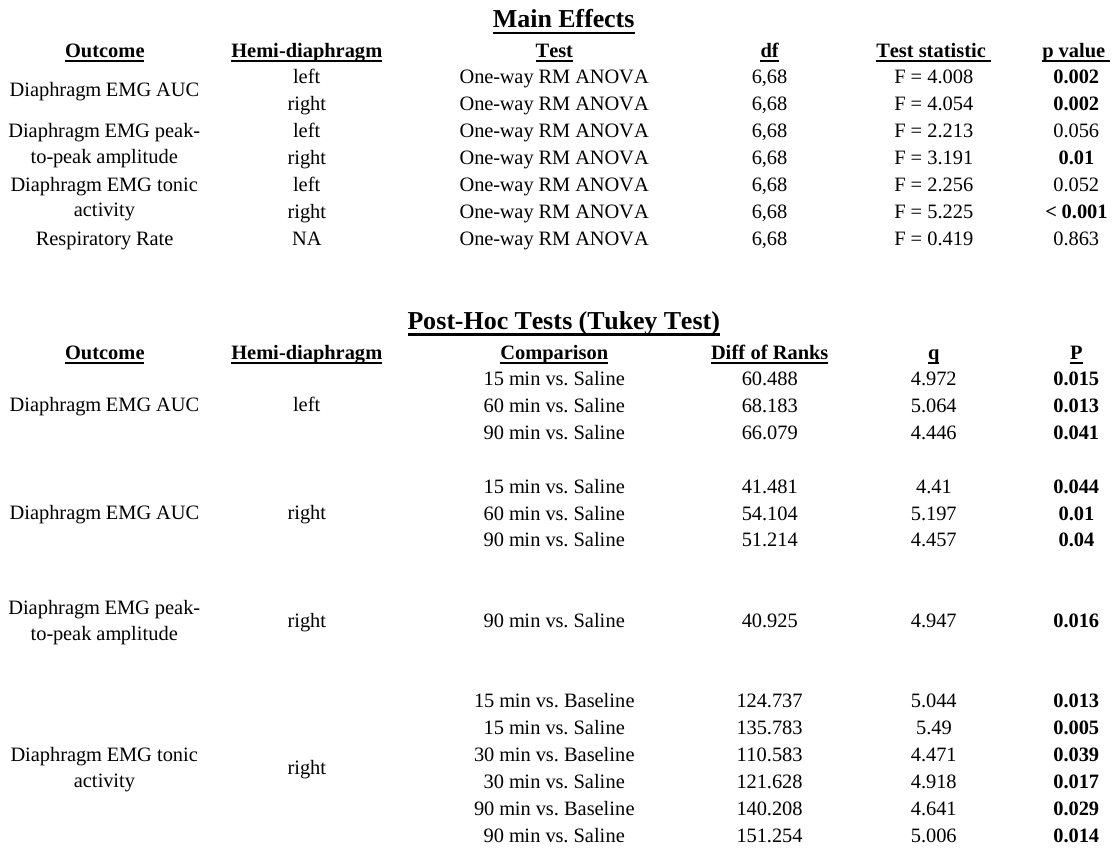

**Supplemental Table 1 (S1). *Statistical summary for the impact of DREADD activation on diaphragm EMG in wild-type mice.*** Time points are in reference to minutes passed since J60 infusion. Summary data is presented in Figure 1. EMG = electromyography, AUC = area under the curve, RM = repeated measures, df = degrees of freedom. Bolded p-values indicate p < 0.05.

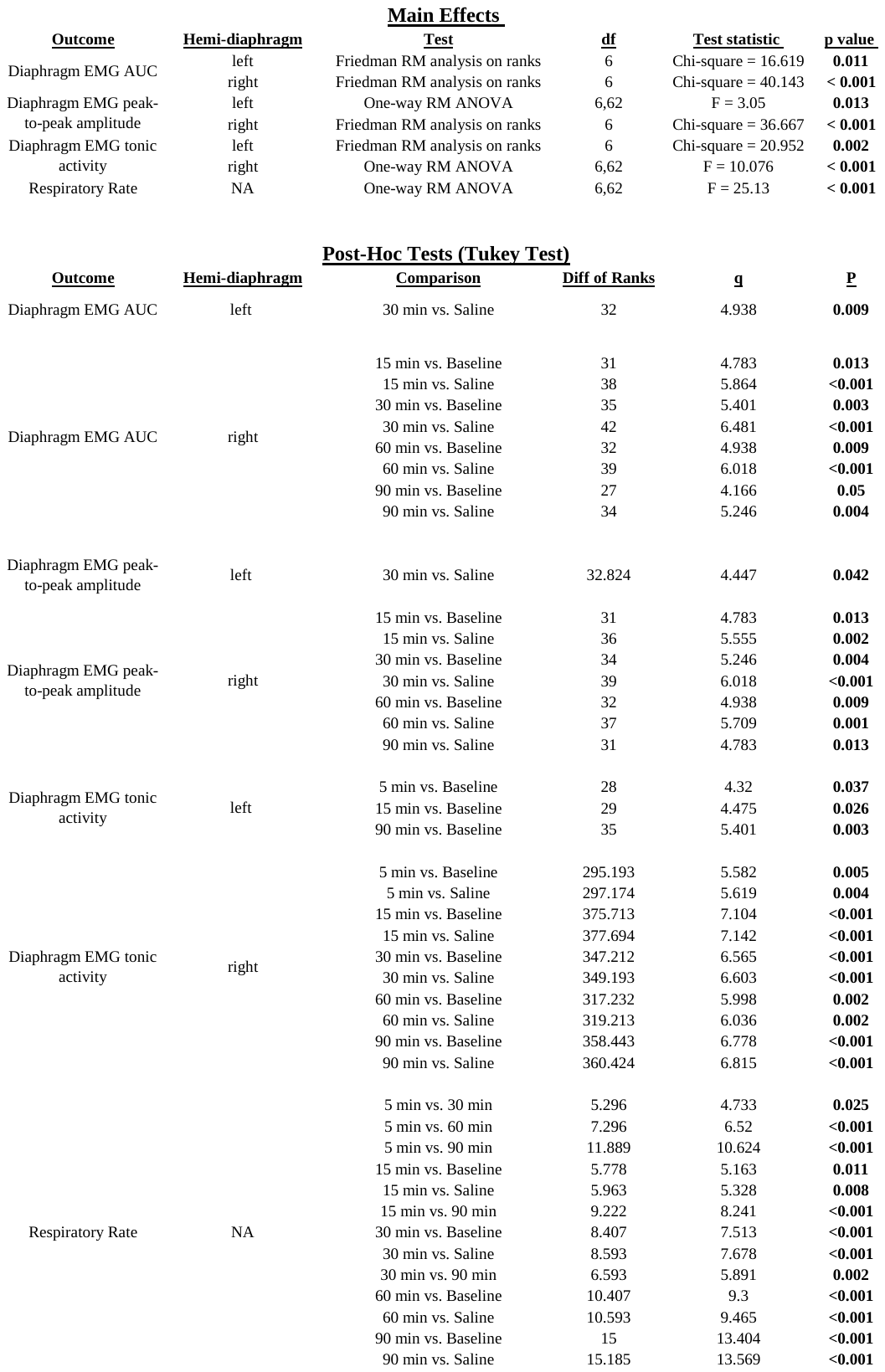

**Supplemental Table 2 (S2). *Statistical summary for the impact of DREADD activation on diaphragm EMG in ChAT-Cre mice.*** Time points are in reference to minutes passed since J60 infusion. Summary data are presented in Figure 2. EMG = electromyography, AUC = area under the curve, RM = repeated measures, df = degrees of freedom. Bolded p-values indicate p < 0.05.

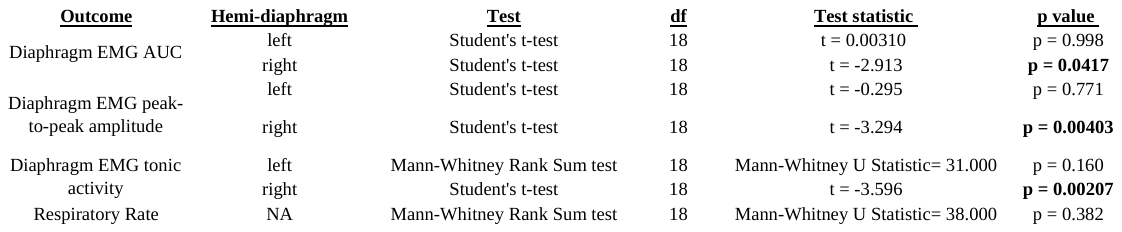

**Supplemental Table 3 (S3). *Statistical summary for the impact of DREADD activation on diaphragm EMG in wild-type mice vs. ChAT-Cre mice at the 30-min post-J60 infusion time point.*** Summary data are presented in Figure 3. EMG = electromyography, AUC = area under the curve, df = degrees of freedom. Bolded p-values indicate p < 0.05.

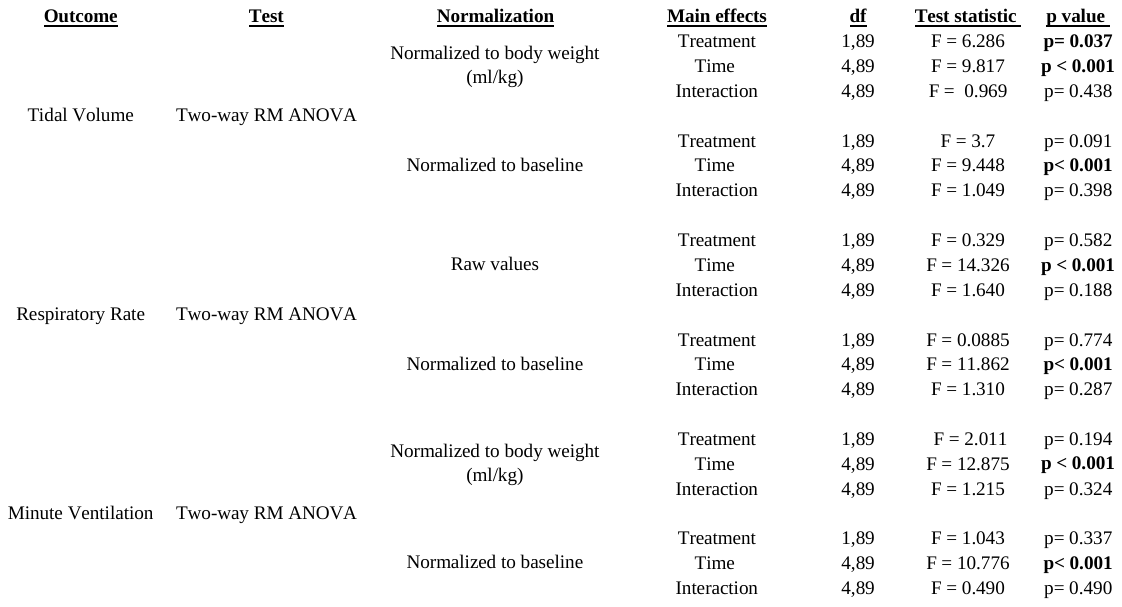

**Supplemental Table 4 (S4). *Statistical summary for the impact of DREADD activation on plethysmography outcomes in unanesthetized ChAT-Cre rats using two-way repeated measures ANOVAs.*** Each outcome measure is presented normalized to body weight (with the exception of respiratory rate) and normalized to values at baseline. Summary data are presented in Figure 4. RM = repeated measures, ml/kg = milliliters of air per kilogram of animal’s body weight, df = degrees of freedom. Bolded p-values indicate p < 0.05.

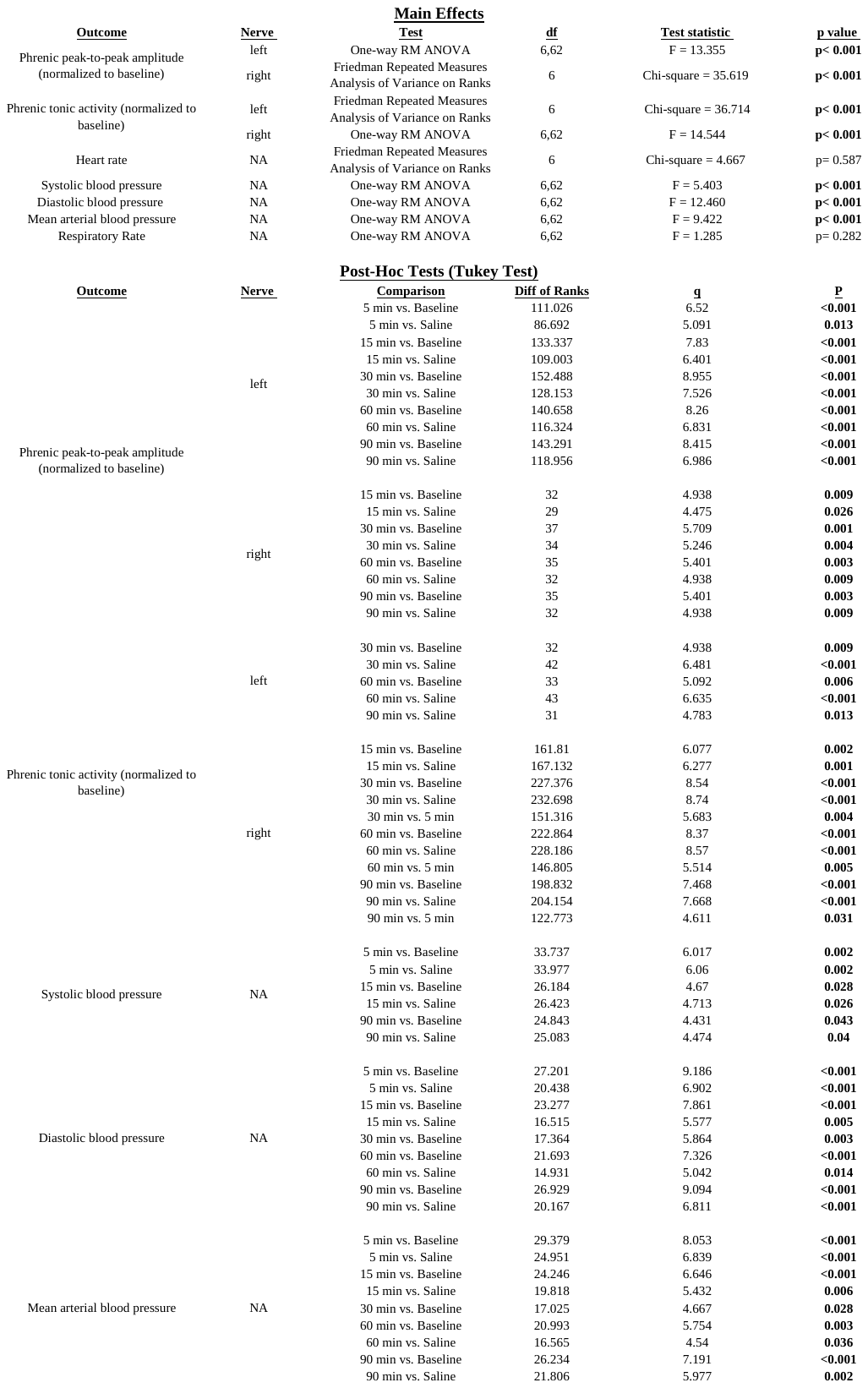

**Supplemental Table 5 (S5). *Statistical summary for the impact of DREADD activation on phrenic nerve activity in ChAT-Cre rats.*** Time points are in reference to minutes passed since J60 infusion. Summary data are presented in Figure 5. RM = repeated measures, df = degrees of freedom. Bolded p-values indicate p < 0.05.
